# *Descriptron*: Testing Artificial Intelligence for Automating Taxonomic Species Descriptions with a User-friendly Software Package

**DOI:** 10.1101/2025.01.07.631758

**Authors:** Alex R. Van Dam, Liliya Štarhová Serbina

## Abstract

Recent advances in Computer Vision, Convolutional Neural Networks (CNNs), Vision Transformers (ViTs) and Large Language Models (LLMs) suggest that it may be possible to approach mimicking the ability to decode knowledge about morphology and taxonomy to describe species in an automated way. Here we test for the first time a current state-of-the-art Vision Language Model (VLM) to approximate taxonomic species descriptions in an automated manner. The test utilizes a new graphical user interface, *Descriptron*, that collects data about biological images and transmits this highly specialized knowledge to a VLM to decode the taxonomic knowledge encoded in labeled biological images with text. Our results indicate that current state-of-the-art VLM (GPT-4o) can produce automated species descriptions that with error correction approximate taxonomists’ ability to describe morphological features of species and organize them in taxonomic keys. However, the results are not without significant errors and the VLM requires the input of the taxonomists knowledge to prevent widespread hallucinations by the VLM. We find here that the taxonomist is clearly needed to both teach and closely supervise the VLM. However, the time saved by utilizing *Descriptron* is also significant. Taxonomists remain essential for teaching and closely supervising the VLM. The time saved by utilizing *Descriptron* are nevertheless already very significant. The *Descriptron* program and supporting example prompt files are free to use under an Apache2 License available from: https://github.com/alexrvandam/Descriptron.

---

The number of species in biological collections waiting to be described varies from dozens to thousands depending on the institution collection size and taxonomic group (Fontaine, Perrard & Bouchet, 2012). Across taxonomic groups undescribed species often languish in collections, taking on average 21 years from discovery to description (Fontaine, Perrard & Bouchet, 2012).

This delay is particularly concerning in light of the ongoing global decline in biodiversity caused by human activities, emphasizing an urgent need for accelerated taxonomic efforts. The most diverse Class of metazoans (Arthropoda) contains more than 1.01 million species described in the scientific literature (Stork, 2018). Multiple data sources point to the total estimated number of arthropods species to be approximately 7 million (Stork, 2018). Because taxonomists describe about 7,000 new species of arthropods a year (Stork, 2018), it will take 480 (Mora et al., 2011) to ∼850 more years to describe the remaining arthropods on Earth. The majority of species are in tropical regions, and many arthropod species will go extinct before taxonomists can describe them (Stork, 2018). Another problem is arthropod taxonomists typically have a backlog of new species awaiting description, so solving this could also help speed up the biodiscovery process (Fontaine, Perrard & Bouchet, 2012). Data collection is a particularly laborious step in taxonomic species descriptions, with a median time of 12 years from discovery to description for many arthropods (Fontaine, Perrard & Bouchet, 2012). This lengthy timeline reflects the need for fine-grained observational data.

Clearly, improvements to the rate of description of new species will increase our understanding of biological diversity of life on Earth. Over the last eight years, a flurry of research has aimed to automate geometric morphometrics (quantifying differences in shape) (Adams, Rohlf & Slice, 2013; Karanovic, Djurakic & Eberhard, 2016; Van Belleghem et al., 2018; Hsiang et al., 2018; Porto & Voje, 2020; Porto, Rolfe & Maga, 2020; Ginot & Debat, 2022; Wiese et al., 2022; Weller et al., 2022; Mitteroecker & Schaefer, 2022; Di Martino et al., 2022). There has also been a large number of papers attempting to use machine learning to automate insect identification over the last five years with hundreds of papers mentioning “machine learning”, “image identification”, and “insect” in their keywords (Gao et al., 2024). There have even been artificial intelligence (AI) classification models that approach expert taxonomist level of accuracy (Valan et al., 2019). But these are only a subset of the information needed in a taxonomic species description. A promising approach to solve this problem is to link morphological characters in a homology based framework via a new computer language *Phenoscript* and *python* library *Phenospy* (Tarasov et al., 2023; Girón et al., 2023; Montanaro & Tarasov, 2024). These languages allow users to describe new species using character based ontology lookup frameworks and automatically compare phenotypes (Tarasov et al., 2023; Girón et al., 2023; Montanaro & Tarasov, 2024). There has also been recent efforts to help make advanced species delimitation software more accessible for taxonomists through a user-friendly graphical user interface (*iTaxoTools*), which is another way to help speed up the process of taxonomic species descriptions (Vences et al., 2021). Species delimitation tools that include artificial intelligence also hold significant promise (Karbstein et al., 2024). Currently, no automation of the collection and subsequent description of finer-grained characters, for taxonomic species descriptions exists. Additionally in some of the most diverse taxonomic groups more than 50% of new taxonomic descriptions are performed by retired taxonomists and self-taught citizen scientists <u>(Fontaine, Fontaine & Prévot, 2021)</u>. Solutions that could compliment and work synergistically with ongoing efforts in geometric morphometrics, image based automatic clade identification, ontology-based computer scripting, artificial intelligence tools and classic taxonomy could be valuable in helping to speed up the description of new species.

Despite a forecast of hundreds of years to describe the remainder of arthropod biodiversity, in the face of anthropogenic driven extinction events <u>(Mora et al., 2011; Outhwaite,</u> <u>McCann & Newbold, 2022)</u>, there is hope to make rapid progress. Globally the same 20 families of flying insects make up greater than 50% of the species diversity regardless of clade age, habitat type, climactic region, or continent (Srivathsan et al., 2023). These 20 clades are also amongst the most neglected in terms of taxonomic descriptions making them ‘Dark Taxa’ (Srivathsan et al., 2023). Advances in rapid parallel barcoding of individual arthropod specimens in a non-destructive way have been developed to feed into a bioinformatic pipeline lending itself to rapid species delimitation and clade placement (Hartop et al., 2022; Srivathsan et al., 2023; Meier et al., 2024). In addition to this advances in robotics have been harnessed to automate the sample collection of arthropods (Wührl et al., 2022) making it possible to begin to speed up the overall discovery process of novel species. The potential for taxonomists to focus on these 20 ‘Dark Taxa’ clades and build custom models and bioinformatic pipelines to describe them would have a significant impact on our understanding of global biodiversity.

Robust computer vision techniques packaged in a user-friendly graphical user interface, may allow ‘Dark Taxa’ and their overlooked aspects of morphology, or what the evo-devo community refers to as the ‘phenome’, to be analyzed by new AI tools, harnessing existing, massive datasets in new ways. This paper describes the very first implementation of a new program “*Descriptron*” (Van Dam, 2024) to automate geometric morphometrics via semi- landmarks, extraction of key morphometrics, quantitative color metrics, overall color pattern and shape of specific user defined morphological features. In addition to that *Descriptron* attempts to go a step further and use these fine-grained morphological details to constrain both encoder only and autoregressive Large Language Models (LLMs) to produce draft taxonomic species descriptions.

To be clear the hypothesis the authors are testing here is if recent advances in computer vision in particular fully convolutional neural networks, can be used to extract contour data from existing images to help constrain and inform vision transformer encoder and autoregressive transformer LLMs to produce coherent draft species descriptions. Currently it is completely unknown if such models can extract fine grained characters from arthropods or the degree of training needed to execute draft species descriptions. Here we plan to test if it is possible to produce partial draft species descriptions, using a small number of specimens from the superfamily Psylloidea (Insecta: Hemiptera: Sternorrhyncha) as a test group. We then compare the output from *Descriptron* to expertly crafted partial species descriptions of the same taxa to see if this approach is plausible.

## Materials and Methods

### A Brief Introduction to Task Relevant Deep Learning Computer Vision Methods

*Summary of relevant deep learning methodologies.*—Deep learning is a subfield of machine learning that is based on the use of artificial neural networks (ANNs) to model and solve complex problems. ANNs are composed of layers of interconnected nodes that process and transform input data in a hierarchical manner (Goodfellow, Bengio & Courville, 2016). Deep learning has shown impressive performance in various applications, including image and speech recognition, natural language processing, and game playing (LeCun & Bengio, 1995; LeCun, Bengio & Hinton, 2015). One of the key advantages of deep learning is its ability to automatically learn hierarchical representations of data, which can be used to extract high-level features from complex input patterns (Bengio, Courville & Vincent, 2013).

Convolutional Neural Networks (CNNs) are a type of ANN that are designed to process and analyze visual imagery. Unlike other types of ANNs, such as Multi-Layer Perceptrons (MLPs) and Recurrent Neural Networks (RNNs), CNNs learn and extract hierarchical features from raw image data through the use of convolutional layers (LeCun & Bengio, 1995). The main difference between CNNs and other ANNs lies in their architecture and how they process input data (Lecun et al., 1998). Unlike MLPs, which are fully connected networks that process each input feature independently, CNNs use local receptive fields and shared weights to extract features from adjacent regions of the input (Fukushima, 1980; Lecun et al., 1998). This makes CNNs particularly well-suited for image analysis tasks, as they are able to preserve the spatial relationships between adjacent pixels in an image (LeCun, Bengio & Hinton, 2015), and consequently are highly effective at fine grained image classification, object detection, and segmentation. Another key advantage of CNNs is the ability to learn features through backpropagation, i.e., to automatically learn and adapt to new data without explicit feature engineering (Krizhevsky, Sutskever & Hinton, 2012, 2017). This characteristic makes feature-learning by CNNs highly flexible and capable of generalizing to a wide range of tasks and data types. One popular CNN-based framework for object detection and segmentation is *Detectron2* (Wu et al., 2019), which is an open-source software library that provides state-of-the-art models for these tasks (Wu et al., 2019).

Transformers and Vision Transformers (ViT), such as the dense prediction transformer (*DPT*) introduced by Ranftl et al. (Ranftl, Bochkovskiy & Koltun, 2021), were originally designed for natural language processing tasks (eg. *ChatGPT*) (Radford et al., 2019) but have also shown great promise for computer vision tasks (Ranftl, Bochkovskiy & Koltun, 2021; Ranftl et al., 2022). Unlike CNNs, Transformers do not use convolutional layers but instead rely on self-attention mechanisms to learn contextual relationships between different parts of the input data (Vaswani et al., 2017; Ranftl, Bochkovskiy & Koltun, 2021; Ranftl et al., 2022). In particular, *DPT* and the Segment Anything Model (SAM) have allowed for precise image segmentation (Ranftl, Bochkovskiy & Koltun, 2021; Kirillov et al., 2023).

*Detectron2* was chosen as the base CNN for image instance segmentation for *Descriptron* pipeline as it is opensource and was built from the ground up for users to add new models to the existing model zoo backbone in *PyTorch* (Wu et al., 2019). For the fine scale extraction and location of features from *Detectron2* to a statistical framework python offers a wide variety of computer vision options including *OpenCV* (also known as *CV2*). *OpenCV* is classic computer vision in *python* and links the higher-level instance segmentation performed by *Detectron2* with a statistical frameworks implemented directly in *OpenCV* or with other *python* libraries such as *NumPy* (Bradski, 2000; Harris et al., 2020a; OpenCV Contributors, 2021). *Detectron2* also runs in either GPU or CPU mode relatively effortlessly allowing the whole system to be accessible to the widest possible audience (Wu et al., 2019).

As an alternative we also implement instance segmentation of sclerites using no specific fine-tuned training for sclerite segmentation using *Segment Anything Model 2* (*SAM2*) (Kirillov et al., 2023; Ravi et al., 2024). In more detail *SAM2* is a transformer-based architecture utilizing a (ViT) meaning that it can process and input image, divide it up into a fixed patch size and then encode those patches as a sequence of embeddings. An embedding is a series of numerical vectors in a multidimensional space that collectively represent the given input such as a word, sentence, image or other modality (Jia et al., 2014; Ranftl, Bochkovskiy & Koltun, 2021; “GPT- 4V(ision) System Card,” 2023; OpenAI et al., 2024). Then each image patch is flattened and projected onto a fixed-dimensional embedding space using a linear projection layer. The image embedding patches are then given spatial context through a positional embedding. Next the ViT processes the embeddings through multiple layers of self-attention and feedforward networks to capture global dependencies and context in the image. A prompt embedding can optionally be added into the same feature space as the image embeddings (prompts can be points, bounding box, masks, and in some cases text) (Kirillov et al., 2023; Ravi et al., 2024). The prompt embeddings are then fused with the image embedding using a cross-attention mechanism. The prompt can serve as a guidance signal to the image instance segmentation mask decoder as an area of interest. A decoder then up samples the hierarchical feature representations from the encoder to match the original image resolution ensuring that the output mask is same resolution as the input image. Finally, the embeddings from the transformer are passed to a lightweight CNN to identify each pixel belonging to the mask (Kirillov et al., 2023; Ravi et al., 2024).

Prompt driven segmentation by *SAM2* is a very different approach compared to what typically involves a significant amount of training to predict a specific category for instance segmentation (Wu et al., 2019; Minaee et al., 2020; Ranftl, Bochkovskiy & Koltun, 2021; Schwartz & Alfaro, 2021; Ginot & Debat, 2022; Ott & Lautenschlager, 2022; Ravi et al., 2024).

Another advancement in machine learning recently is that of transformer based Large Language Models (LLMs) used to interpret text are paired with ViTs, this class of transformer is commonly referred to as Vision-Language Models (VLMs) used in part for Visual Question and Answer (VQA) tasks (“GPT-4V(ision) System Card,” 2023). At the core of VLMs is the combination of vision models and advanced language models. Each modality is processed separately to produce independent embeddings, before the embeddings from a ViT for example are to be fused with the word token embeddings form the language input (Radford et al., 2021; “GPT-4V(ision) System Card,” 2023). The language embeddings are produced through a process of tokenization where each word or sub-word or even individual character are given a numerical representation based on the individual LLM’s method of tokenization vocabulary (eg. Byte Pair Encoding or WordPiece) (Gage, 1994; Devlin et al., 2019). A special token is usually reserved for the beginning and end of a sentence as well as for where the user places the image embedding in the case of a VQA. Multimodal fusion of embeddings from the ViT and LLM can then be accomplished by a variety of methods such as cross-attention mechanisms or concatenation this allows for integration of information across modalities (Radford et al., 2021; Chen et al., 2024). Currently there are a variety of different VLMs some of which have VQA capabilities, perhaps the models best suited for this task are those involved in text generation given an image and a question of that image as part of a prompt.

One such model is *GPT4-Vision* (*GPT-4V*) that has advanced VQA capabilities (“GPT- 4V(ision) System Card,” 2023). *GPT-4V* is an autoregressive LLM meaning that it generates outputs by predicting the next token in a sequence based on the preceding context using a specialized embedding decoder (“GPT-4V(ision) System Card,” 2023). By effectively computing attention scores between modalities (eg. images and text) *GPT-4V* can effectively align the relevant parts of both the image and the text to decode out the most likely response (“GPT- 4V(ision) System Card,” 2023). In slightly more detail in a VQA scenario the image embeddings are processed by the image encoder and these embeddings can be thought of as keys and values representing the spatial and feature-level details of the image. The embeddings will allow the model to “look up” the relevant visual information after fusion. The text embedding come in the form of the question prompt, they determine which parts of the image (keys and values) the model should focus on to answer the question (“GPT-4V(ision) System Card,” 2023). So carefully prompting in specific data into the question is critical to achieve the desired response given the image. Additionally regions of interest can be highlighted or annotated to help indicate which regions are of the most importance (“GPT-4V(ision) System Card,” 2023). In this cross-attention mechanism, the question drives the search by asking what part of the image is relevant and the image provides details needed to respond.

With that general background now established the authors will next explain the overall strategy followed by a step-by-step explanation of our materials and methods and the code involved in *Descriptron*.

*Overall approach to automate species descriptions.*—LLMs and VLMs and even GPT4 are very prone to “hallucinations” or simply making up information that is not present in the image or completely irrelevant to the query (OpenAI et al., 2024). The main approach to constrain this behavior is to provide expertly crafted prompts from taxonomists and supplement the images with information using instance segmentation from CNNs and measurements from classic computer vision, thus providing a fusion of CNNs and ViTs. To help facilitate this and reduce hallucinations we plan to use instance segmentation to highlight key entomological terms to help direct the attention mechanism of the VLM. In addition to this we plan to concatenate in accurate 2D and optionally 3D morphometric data and color data. To cover the materials examined we use the capabilities of *GPT-4V* to perform optical character recognition (OCR) and natural language processing (NLP) to help translate insect labels. In addition to this we use the general reasoning capabilities of *GPT4* to help automate the production of taxonomic keys. By applying classic computer vision, CNN driven instance segmentation, transformer based *GPT-4V* to provide text output based on annotated images, we plan to provide the community a method to rapidly describe new species.

### The Descriptron Graphical User Interface (GUI)

*Overview of the Descriptron GUI*—*Descriptron v0.1.0*, is an advanced GUI crafted in *python* using *tkinter* to assist in a wide range of computational tasks central to taxonomy and systematics (Python Software Foundation, 2024a). The primary goal is to provide a workflow to help automate taxonomic species descriptions (Figure 1). *Descriptron* supports additional workflows for tasks like image processing, advanced color segmentation, semi-landmarking, machine learning model fine-tuning, geometric morphometrics, morphometrics in 2D or 3D (experimental) and diverse data format conversions through initial analyses (Figure 2). Through user-friendly dialog boxes and structured forms, *Descriptron* collects and validates inputs efficiently, ensuring robust error handling and reducing the risk of mistakes.

**FIGURE 1.**
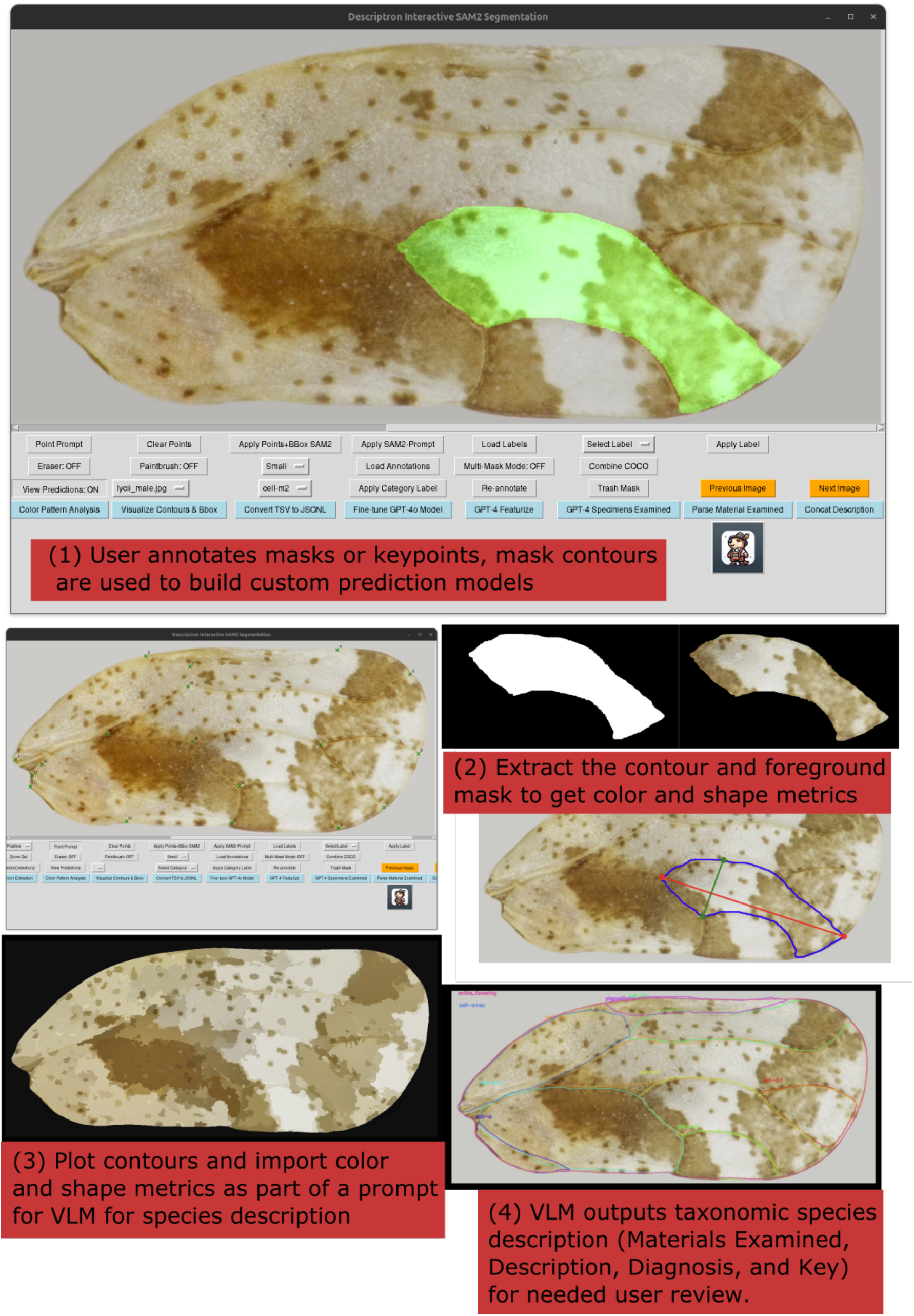
Overall workflow to produce automated taxonomic species descriptions using *Descriptron* graphical user interface.

**FIGURE 2.**
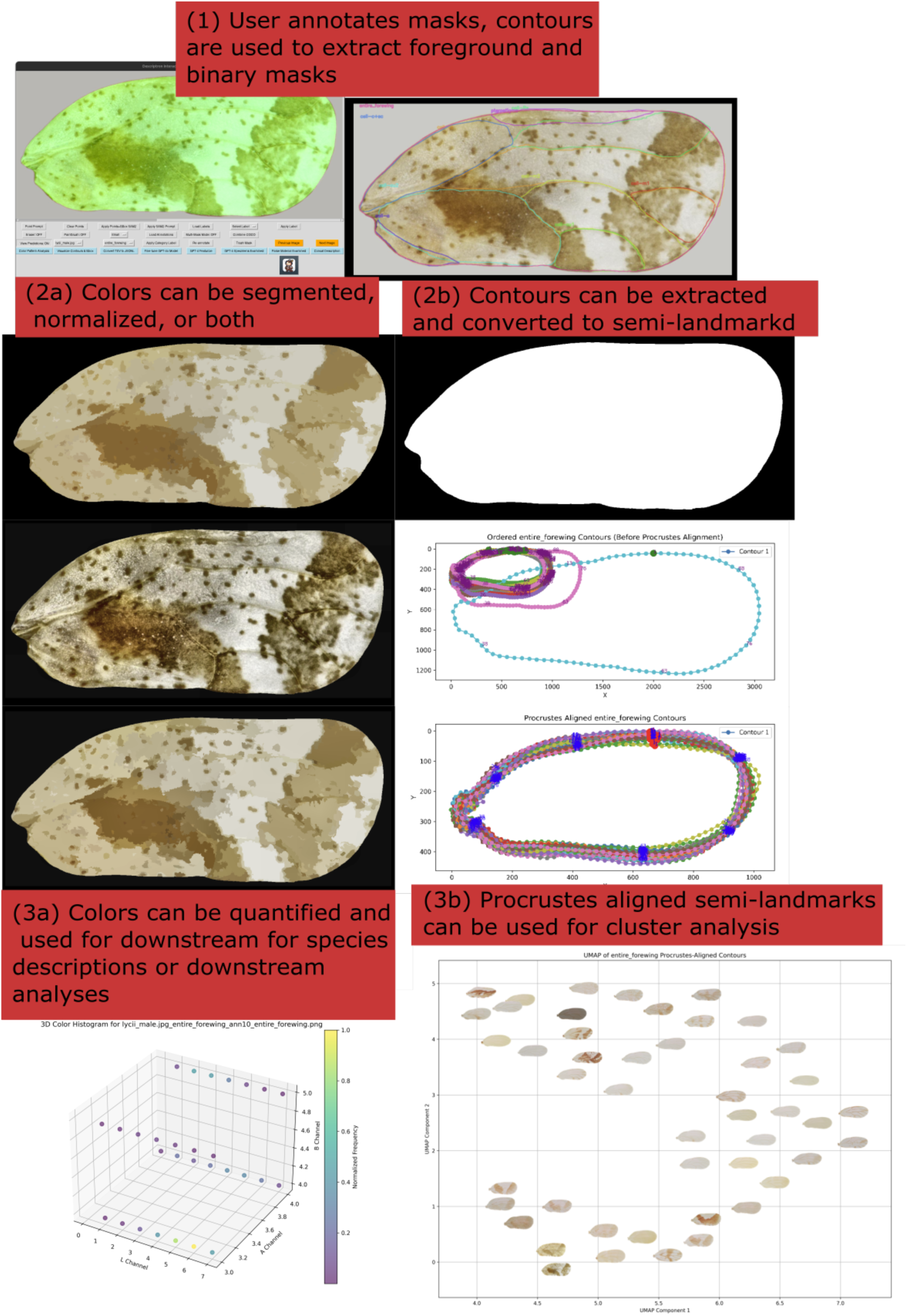
Workflows available within *Descriptron* for (a) quantifying color and color patterns and for (b) automated geometric-morphometrics.

*Environment management and script execution*. —The *Descriptron* GUI employs the *conda run* command to execute scripts within isolated environments, ensuring compatibility and reproducibility (Anaconda, 2024; Anaconda, Inc., 2024). Each module provides clearly defined input fields for selecting files, setting parameters, and specifying output configurations. Input validation is conducted automatically, and scripts are executed in separate threads using the *threading* library to maintain a responsive user interface (Python Software Foundation, 2024b). The *subprocess* module handles the management and execution of these scripts asynchronously, enabling real-time progress indicators, success notifications, and detailed error messages (Python Software Foundation, 2024c).

*Descriptron image processing features.*—*Descriptron* offers tools for generating depth maps, visualizing contours, and overlaying segmentation masks to support morphological and geometric morphometric analyses. The bounding box overlay and keypoint features facilitates precise annotation of regions of interest, which is invaluable for comparative studies of biological specimens and training models for instance segmentation. The image processing functionalities rely heavily on *numpy* for numerical computations and linear algebra operations (Harris et al., 2020b,a). For example, drawing masks from keypoints involves transformations of coordinate systems and matrix manipulations. Similarly, *pandas* is utilized for managing datasets, cleaning data, and aggregating results, ensuring efficient handling of experimental outputs (McKinney et al., 2010). These operations are enhanced by modules like *json* for structured data handling and *sys* for robust system operations (“GPT-4V(ision) System Card,” 2023; Anaconda, Inc., 2024; Python Software Foundation, 2024c,d). By modifying JSON files and updating the viewer on the fly the user gets real-time updates on their annotation workflows by modifying existing files or creating new ones.

### Automatic Instance Segmentation with SAM2

*Incorporating different modes for running SAM2.*—One of the most time-consuming tasks in constructing models for instances segmentation is developing a training data set for the model of choice. The *SAM2* transformer can perform instance segmentation without any user prompts. The masks produced via *SAM2* can then be labeled or thrown out. Taking advantage of this approach *Descriptron* has a workflow available where the user can use the prompt free feature using *SAM2* with default settings that attempt to segment as many details as possible (Ravi et al., 2024). The unlabeled masks can be saved as a COCO JSON (Common Objects in Context (COCO))file and then loaded for later use. Additionally unlabeled masks can be optionally thrown out via the ‘Trash Mask’ button and a simple user provided tab separated list of labels can be loaded via the ‘Load Labels’ button and for on-the-fly mask labeling as well as a custom option for the user to manually type in a new category via the ‘Select Labels’ button. The ‘Apply Labels’ button saves any new additions to the masks that are currently loaded. Finally, the updated masks can be saved and closed using the ‘Marmot Button’ which serves as the method to save and close as well as the *Descriptron* mascot.

*SAM2 prompt and manual segmentation mode.*— The bounding box and point prompts, both negative and positive modes, have been added into the *Descriptron* GUI to work seamlessly to deliver prompts into the *SAM2* workflow. This ability allows for custom generation of specific masks for labeling as before, and takes advantage of *SAM2*’s ability to quickly deliver a custom mask (Kirillov et al., 2023; Ravi et al., 2024). Masks can also be edited with the ‘Paintbrush’ and ‘Eraser’ tools by simply adding or removing pixels to the currently loaded mask. Sometimes it’s desirable to use polygons to draw masks, this can be accomplished by selecting the ‘Positive’ point prompt and then drawing around the desired region. A custom *python* script was written to then convert the keypoints into a mask for downstream editing (labeling, erasing etc.) by the ‘Keypoints to Mask’ button. Keypoints or landmarks can also be saved to COCO JSON format as well using the positive point prompt followed by the use of the ‘Marmot’ button to save the desired keypoints for a given image are placed in their desired locations. The keypoints positions can also have their location moved (but not their numerical order) via the ‘Keypoint Edit: (ON/OFF)’ toggle switch. This allows users to make detailed adjustments to their keypoint annotation workflows.

*File format conversion with Descriptron*.—While not the most intellectually stimulating topic data carpentry can be one of the rate limiting steps in any workflow and deserves more attention (Attwood et al., 2019). Formatting output and file format conversion is found throughout the various buttons and script definitions in *Descriptron*. The ‘COCO ->MinCOCO’ button is specifically for re-formatting standard COCO JSON format to the MinCOCO JSON that *Detectron2* uses (Lin et al., 2014; Wu et al., 2019; McMurray, 2024; Python Software Foundation, 2024d). The COCO JSON format is a widely-used annotation format for object detection and segmentation tasks, designed to be both machine-readable and interpretable by humans (Lin et al., 2014; McMurray, 2024; Python Software Foundation, 2024d). It organizes annotations using hierarchical keys such as images, annotations, and categories, allowing flexibility for bounding boxes, segmentation masks, and object classes. There are a few common variations: (1) Standard COCO JSON: This format keeps all annotations in a single file using nested structures, with clearly designated sections for images, categories, and annotations. (2) COCO JSONL: In JSON Lines (JSONL) format, each JSON object (e.g., representing one image or annotation) is written on a separate line. This format is more suitable for streaming and large datasets. (3) MinCOCO JSONL: A specialized compact format where all data is concatenated into a single line, like a single-line FASTA file. This structure is more efficient for specific workflows but is less human-readable (McMurray, 2024).

Even though JSON is like XML format which users familiar with evolutionary biology software eg. BEAST (Drummond & Rambaut, 2007), in its hierarchical structure—using keys, brackets, and indentation to represent relationships and nested categories—JSON is generally preferred for machine learning workflows due to its lightweight syntax and compatibility with modern programming tools eg. *pytorch* (Paszke et al., 2019). Knowing the distinctions between these variants can greatly expedite troubleshooting and processing. *Descriptron* simplifies these tasks by ensuring data is output in formats compatible with popular frameworks and downstream processes.

### Instance Segmentation Descriptron Workflow with Detectron2

*Conversion to COCO format.*—*Detectron2* is a robust framework for object detection and segmentation built by Facebook AI Research (FAIR) and is particularly well-suited for handling large-scale, annotated datasets. Below, are the steps involved in training a Mask R-CNN model using *Detectron2* for instance segmentation.

A custom function convert_to_coco converts annotations to COCO format (Lin et al., 2014), ensuring each sclerite category is assigned a unique class ID. This involved extracting polygon coordinates for each sclerite and calculating bounding boxes. The COCO format is a standard for object detection and segmentation datasets and allows for easy integration with the *Pytorch* and *Detectron2* framework (Paszke et al., 2019; Wu et al., 2019).

*Model configuration and training.*—The authors used a transfer learning approach by fine tuning the existing Mask R-CNN model (He et al., 2017). First a configuration object can be modified to suit specific training needs is loaded. Next a configuration object with settings from a predefined YAML file. The YAML file contains configurations for training a Mask R-CNN model with a ResNet-50-FPN backbone on the COCO dataset (He et al., 2015; Lin et al., 2017). The model_zoo.get_config_file function retrieves the appropriate configuration file from the *Detectron2* model zoo. All user specified classes can be trained at the same time so that the model can learn about the new classes together in context of one another. The backbone build_resnet_fpn_backbone combines a Residual Nerual Network (ResNet) with a Feature Pyramid Network (FPN), providing rich feature hierarchies for object detection (He et al., 2015; Lin et al., 2017). Key configurations included setting the batch size to 2 images per batch, in other words two images get processed before updating the model parameters. Larger batch sizes can lead to faster training per epoch but require more memory, while smaller batch sizes can make the model updates noisier but often lead to better generalization.

*Learning rate and model training iterations.*—The training script employs a warmup phase of 3,000 by default but users can specify their own number of warmup iterations with an initial learning rate set to 0.0001, a slow but accurate learning rate. After the warmup phase, the learning rate is dynamically adjusted based on the number of GPUs. Specifically, the base learning rate is set to 0.00025 multiplied by the number of images per batch. For instance, if there are two GPUs, each processing 2 images per batch, the effective learning rate becomes 0.00025 * 2 = 0.0005. The learning rate is further scheduled to decrease at specific iterations: it will drop by a factor of 10 at 40% of iterations and again at 80% of iterations, helping the model to converge by reducing the learning rate as training progresses. The total training process is set to run for a maximum of 35,000 iterations by default but can be user defined. This dynamic adjustment and scheduled decrease in learning rate aim to balance the need for precise learning and the risk of overfitting. While this approach helps to prevent overfitting, it may result in slower learning. Future implementations will explore and define an optimal learning rate specifically for arthropod sclerites, but for now, we use this adaptive approach.

*Regions of interest per image.*—The learning rate is influenced by the number of Regions of Interest (RoIs) sampled per image during training. In this script, the RoI batch size per image is set to 128, which is a balanced number designed to prevent crashing a single GPU or CPU. RoIs are candidate regions proposed by the Region Proposal Network (RPN) that are used to compute the loss and update the model weights (Wu et al., 2019). Increasing the number of RoI batch size considered for each image can enhance the model’s ability to learn from a greater

variety of examples within each image, thereby improving performance. However, a higher RoI batch size necessitates more memory resources and possible memory crash.

*Number of images processed per batch.*—The total effective batch size for gradient computation is determined by the number of GPUs and the number of images processed per batch. This, in turn, affects the overall learning rate. The number of GPUs used not only speeds up the training process but also influences the learning rate by scaling it according to the batch size, which can impact the rate of learning and the risk of overfitting. Thus, the configuration and number of GPUs play a critical role in determining both the efficiency and effectiveness of the training process (Wu et al., 2019).

*Data augmentation.*—To enhance model robustness, various data augmentation techniques were applied such as random rotation, brightness, contrast, and saturation adjustments. These augmentations help the model generalize better by exposing it to varied input data during training using the built in *Pytorch* and *Detectron2* libraries (Paszke et al., 2019; Wu et al., 2019). A custom trainer class, AugmentedTrainer, was used to incorporate these augmentations and handle the training process.

*Early overfitting halt.*—A custom early stopping function was implemented using an EarlyStoppingHook based on validation performance to prevent overfitting if segmentation average precision scores above 50% (AP50) did not improve after 5,000 iterations. AP50 specifically refers to the Average Precision calculated at as the ratio of area of overlap divided by area of union (IoU) of the predicted segmentation mask with a threshold of 0.50. AP50 can be used as a measure of accuracy for both bounding box (class prediction) and instance segmentation (how well the prediction matches the ground truth) (Everingham et al., 2010). This means that a predicted bounding box (or segmentation mask) is considered a true positive if its IoU with the ground truth box (or mask) is greater than or equal to 0.50.

*Weight decay.*—This parameter applies a regularization term to the loss function to prevent overfitting. Weight decay penalizes large weights in the model, encouraging the model to keep the weights small and simple (Wu et al., 2019). The parameter of 0.0001 was set for weight decay, as this helps improve generalization by adding a small penalty proportional to the magnitude of the weights.

*Gradient clipping.*—Gradient clipping is a technique to prevent the gradients from exploding during the backpropagation process. The clip type was set to a value of 1.0. A gradient value type of 1.0 means that any gradient value greater than 1.0 will be scaled down to 1.0 this is done to prevent instability in the training process.

*Model region of interest heads.*—This parameter sets the number of output classes for the model, specifically for the Region of Interest (RoI) heads. This was designed to adjust dynamically based on the number of classes in the novel training data allowing a user to only specify the class names in their training data to set this parameter. The advantage of this is to ensure that the model’s output layer matches the number of categories in the dataset. This is crucial for accurate classification and segmentation, as the model needs to correctly identify and differentiate between all the different categories.

*Training loss visualization.*—A custom hook LossPlotHook was developed to visualize the training loss curves. This hook plots the loss every 20 iterations and saves the plot as a PNG file, allowing for monitoring of the training process and ensuring the model is learning appropriately.

*Evaluation and saving the model*. —After training, the model was evaluated using *COCOEvaluator* on the validation dataset (Paszke et al., 2019; Wu et al., 2019). This evaluator calculates metrics such as average precision (AP) and average recall (AR) to assess the model’s performance. A custom plot script was written to dynamically plot these statistics as they become available every 1,000 generations but can be set by an input argparse statement. The manual evaluation of this plot is critical to assess the potential of overfitting. If the loss plot continues to decrease but the AP and AP50 scores plateau or show stochasticity, then overfitting is more of a risk. Generally, it is best to stop the model shortly after a AP50 plateau is reached.

*Saving the model, evaluation statistics, and configurations.*—The trained model, configuration files, and metadata were saved for future use. This is critical for retraining the model with the novel sclerite classes. This allows for other users to plugin new training data using the same models and or update with new training classes. The results of the evaluation, including the AP and AR metrics, were summarized and saved in a JSON file every 1,000 generations (or via user specified input) as the model updates.

*Novel Fine Tuned Model Prediction on Test Data Set with Detectron2 Prediction model settings.*—The prediction uses *python 3.8*, *pandas*, *CV2*, *numpy*, *json*, *Pytorch* and *Detectron2* codebase (Bradski, 2000; Foundation, 2019; Paszke et al., 2019; Wu et al., 2019; Harris et al., 2020a; OpenCV Contributors, 2021; Anaconda, 2024; McMurray, 2024). First the default model is called so that the different parameters can be set. During training a configuration YAML file is saved with the model parameters used, this file is loaded next to set hyperparameters and model architecture. Next the parameter that specifies valid predictions was set with a confidence score above 0.5, meaning that predicted detections below this confidence threshold will be filtered out.

Next the training model weights from the fine-tuned *Detectron2* model are loaded. Model weights are used to initialize the model for inference, providing a starting point that has been fine-tuned on the user specified training data. The learning rate is explicitly set to 0.0001 by default or a user specified value and the maximum number of iterations can also be set.

Similarly, the number of model RoI heads and classes was set to the same number as in the training step by default. The class number is set dynamically based on the metadata JSON file created during the training process. This also ensures that the predicted classes names will match with the desired output. Many of the same parameters are repeated in the training to help ensure that the configuration is complete and ready for any potential further training. An evaluation script is provided but this part can be ignored if the prediction script is used on novel test data for inference only.

### Automated Semi-Landmarking of contour data for automated 2D morphometrics and geometric morphometrics

*Contour extraction.*—Taking the sclerite contours from the previous step, in order to automate the process of geometric morphometrics analysis by detecting, normalizing, and aligning contours from a set of images another novel script needed to be written. The script leverages several advanced image processing and machine learning libraries to ensure precise alignment and comprehensive analysis of the contours. The workflow involves resampling contour points to a uniform number, aligning contours using Principal Component Analyses or ‘PCA-based’ methods, performing Procrustes transformation and analysis for shape comparison, and saving the results in various formats for further examination and visualization. It also leverages the ability to describe all of the summary measurements about the contour, such as length width, area etc. To achieve these tasks, the script incorporates multiple key modules, including *CV2*, *NumPy*, *SciPy*, *scikit-learn*, and *Matplotlib* (Bradski, 2000; Hunter, 2007; Tosi, 2009; Pedregosa et al., 2011; Van der Walt et al., 2014; AI, 2020; Virtanen et al., 2020; Harris et al., 2020a; OpenCV Contributors, 2021; Google, 2024; Jones et al., 2024; Smith, 2024).

A custom python script uses the predicted contours to extract their coordinates from either all predicted classes within a folder or a specific class (eg. pterostigma) from a given folder. The contours are then converted into binary masks, foreground masks with the RGB color data, NumPy array files (npy), simple xy coordinates text files and TPS files. The binary masks can also be used as training data for a variety of different machine learning algorithms. The number of instances found per image is recorded and the range across a given list of samples can be specified.

*Automated conversion of binary and foreground to automated semi-landmarks and NumPy arrays for semi-landmarking.*—*OpenCV* is used for essential image processing functions such as color conversion, contour detection, and geometric transformations (OpenCV Contributors, 2021). *NumPy* provides the foundational array operations necessary for handling and manipulating the contour data (Harris et al., 2020b,a). *SciPy*’s interpolation and Procrustes analysis functionalities are crucial for resampling contour points and performing shape analysis (Virtanen et al., 2020). Via *Scikit-learn* a Principal Component Analyses (PCA) orients the samples and scaling tools are employed for contour alignment and normalization (Pedregosa et al., 2011).

The script includes several custom functions that execute specific tasks in the workflow. Functions like find_lowest_rightmost_point and order_points_clockwise ensure that the contour points are ordered consistently. The resample_contour_points_fixed_first function resamples the contour points to a fixed number, facilitating uniformity in subsequent analyses. This last step is very important, because this sets the first point as a single landmark often needed in downstream analyses and the rest of the points as semi-landmarks as they are spaced evenly at uniform distances and by a given user specified number along the contour. The last step essentially sets the homology for the single keypoint or landmark, i.e. the starting position, and the semi-landmarks for downstream geometric morphometric analyses. The align_contours_to_reference function aligns contours to a reference using PCA-based orientation, while procrustes_analysis performs shape alignment and comparison. Connections between the points are drawn using linear interpolation implemented via *numpy* (Harris et al., 2020b), which is the simplest method, connecting two consecutive points with a straight line. It calculates intermediate points by assuming a constant rate of change between known data points (Harris et al., 2020a). While straightforward and computationally efficient, linear interpolation can result in sharp angles at each data point, leading to a piecewise linear approximation that may not capture the smoothness of natural curves. What this means is the number of semi-landmarks should be sufficient in order to retain the contour shape information (Mitteroecker & Gunz, 2013; Watanabe, 2018).

*Matplotlib* is utilized for visualizing the contours and their transformations (Hunter, 2007; Tosi, 2009).

The script also includes higher-level functions for processing entire folders of images and contours. The process_image function handles individual image processing, extracting scale bar lengths and text information. The process_folder function iterates over all images in a specified folder, applying process_image and saving the results. Similarly, process_contours reads contour files, calculates various metrics, and saves these metrics to text files. The resample_and_save_contours function orchestrates the resampling, alignment, and Procrustes analysis of contours, saving the processed data in formats such as CSV, TPS, and NPY files.

*Contour visualization.*—Visualization functions like visualize_contours are provided to plot and inspect the contours visually. Visual outputs are generated to illustrate measurements and contours. Using Matplotlib and OpenCV, annotated images display object contours as well as the transformed semi-landmarked contours.

*Clustering and dimensionality reduction*.— PCA reduces feature dimensions to 2D components for linear analysis (Virtanen et al., 2020). UMAP (via *umap-learn*) provides non-linear dimensionality reduction for clustering (McInnes, Healy & Melville, 2018; Wang et al., 2021). Scatterplots overlay thumbnails of segmented objects at their respective PCA/UMAP coordinates using OffsetImage from *Matplotlib*. Clustering is performed using DBSCAN (Density-Based Spatial Clustering of Applications with Noise) and Hierarchical Clustering (via *SciPy*’s linkage and dendrogram functions). DBSCAN identifies clusters while treating noise as outliers, and hierarchical clustering produces dendrograms for visualizing group relationships.

*2D segmentation mask morphometrics.*—*EasyOCR* and *OpenCV* enables text extraction from images, which is used for detecting scale bars and their units if present in the original image (Bradski, 2000; AI, 2020; OpenCV Contributors, 2021). The ‘Run Measurement Script’ button functions and definitions detect and measure the scale bar length in images using *opencv*, using color thresholding and then are passed for OCR and NLP analyses using *EasyOCR* (AI, 2020).

EasyOCR employs a hybrid architecture that combines CNNs and Recurrent Neural Networks (RNNs) for its text recognition tasks. Specifically, it utilizes the CRNN (Convolutional Recurrent Neural Network) model, which integrates CNNs for feature extraction and RNNs for sequence modeling, followed by a Connectionist Temporal Classification (CTC) decoder for transcription (AI, 2020). This scale information is vital for converting pixel measurements to physical units like micrometers. Using *OpenCV* the contrast between the background and the scale bar is used to locate it as a thin geometric horizontal line with high contrast compared to the background.

This is then used to convert from the unit of measurement, usually in mm for most specimen photos to pixels as well as to μm if “mm” is detected by *EasyOCR*.

A custom definition was written to calculate the maximum length and height of a given instance segmentation mask by the using *opencv* and *sklearn* PCA (Bradski, 2000; Pedregosa et al., 2011; Van der Walt et al., 2014; OpenCV Contributors, 2021), which is based on the shape and not the orientation of the mask. The principal component is taken as the length with the width being calculated as the maximum internal line within the mask contour that is perpendicular to the principal component. This method may be erroneous in some cases to very odd shapes that are wider than they are long but for most categories this measurement is orientation invariant allowing it to be used on a wide variety of images. The area, perimeter, hull, hull area, solidity, and equivalent diameter are all calculated from the contour directly using *opencv* (OpenCV Contributors, 2021).

*Data aggregation and output.*—Morphometric measurements, including length, width, and corresponding coordinates, are saved as individual text files, a single CSV file, or a JSONL file. Instance counts and group-level statistics, such as mean, standard deviation, and range, are computed using Pandas.

### Preliminary 3D Morphometrics

*Camera intrinsics determination*.—Annotations were processed to extract relevant object contours necessary for subsequent metric calculations. The images were stored in a designated directory, ensuring that only images present in the specified directory were included in the analysis to maintain consistency and relevance.

Accurate metric measurements in real-world units require precise camera intrinsic parameters, specifically the focal lengths (𝑓_𝑥_, 𝑓_𝑦_) and the principal points (𝑐_𝑥_, 𝑐_𝑦_). The initial approach involved extracting EXIF metadata from each image to retrieve camera specifications, including make, model, focal length, sensor dimensions, and if available pixel sensor size (OpenCV Contributors, 2021). Focal lengths provided as rational numbers in the EXIF data were correctly converted to floating-point values to ensure unit consistency. The focal length in pixels was calculated using the pinhole camera model equations:

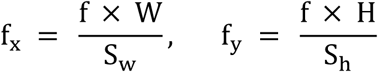

where 𝑓 is the focal length in millimeters, 𝑊 and 𝐻 are the image width and height in pixels, S_w_ and S_h_ are the sensor width and height in millimeters, respectively (OpenCV Contributors, 2021).

For images lacking comprehensive EXIF data, a scale bar detection algorithm was employed as in the the 2D measurement methods previously outlined in this paper. EXIF data can be retrieved via *pillow* (Umesh, 2012; Clark, 2015). The scale bar’s length in pixels (L_pixels_) was measured using image processing techniques, and its real-world length in millimeters (𝐿_𝑚𝑚_) was determined via Optical Character Recognition (OCR) using EasyOCR as previously outlined in the 2D measurement section (AI, 2020). The pixels-per-millimeter ratio (ppmm) was calculated as:

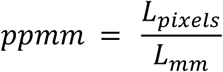

This ratio facilitated the estimation of camera intrinsics based on known sensor dimensions. If both EXIF data extraction and scale bar detection were unsuccessful, users were required to provide intrinsic parameters through command-line arguments, ensuring that the analysis proceeded with accurate and reliable data.

*Depth map generation and calibration*.—Depth maps were generated using the Metric3Dv2 model (Hu et al., 2024a,b), a deep learning framework specialized for monocular depth estimation. The model was pretrained on diverse datasets and fine-tuned for the specific application. Depth maps can be calibrated via Metric3D using camera intrinsics (Yin et al., 2023; Hu et al., 2024b). If camera intrinsics are partially complete to ensure metric accuracy, depth maps are provided can be generated with Metric3Dv2 (Hu et al., 2024a). However plotting the points using back projection still needs to be solved we outline one possible solution below.

*3D measurements using back projection and principal component analysis (PCA)*—Object contours extracted from segmentation masks were back-projected into 3D space using the calibrated depth maps and camera intrinsics. Metric3D does not directly provide code for metrology for contour data, so we had to write our solution to take measurements using this forward-facing 3D data. The back-projection process converted 2D image coordinates (𝑢, 𝑣) to 3D real-world coordinates (𝑋, 𝑌, 𝑍) using the following equations:

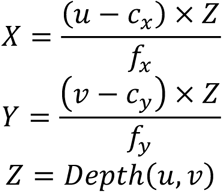

where 𝐷𝑒𝑝𝑡ℎ(𝑢, 𝑣) is the depth value obtained from the depth map at pixel coordinates (𝑢, 𝑣). Principal Component Analysis (PCA) was then applied to the set of 3D points to determine the principal axes of each object. The primary principal component aligned with the object’s length, and the secondary component, perpendicular to the primary, represented the object’s width. The length (𝐿) and width (𝑊) were computed as the maximum distances along these principal axes:

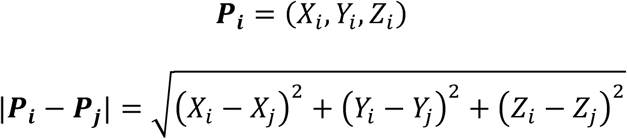

where 𝑃_𝑖_ 𝑎𝑛𝑑 𝑃_j_ are points along the principal component, and 𝑃_𝑘_𝑎𝑛𝑑𝑃_𝑙_ are points along the secondary component. The width was then calculated as the longest interior line along the secondary component that is perpendicular to the principal component. The distance between points could then be defined by the following formulas:

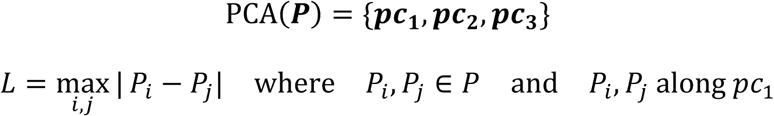

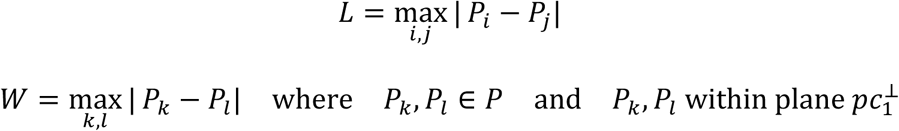

The basic width calculation then has to be plugged into the back projection of the 2D to 3D points as

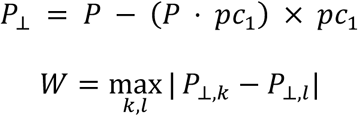

By projecting points onto a plane perpendicular to 𝒑𝒄_𝟏_the width measurement is inherently orthogonal to the object’s length, ensuring that length and width are independent dimensions. Calculating the maximum distance within this plane ensures that the width reflects the object’s internal breadth rather than external bounding constraints (eg. bounding box), providing a more accurate and meaningful metric invariant to region of interest or image orientation.

*Clustering and dimensionality reduction*.—To analyze the distribution and clustering of object metrics, various dimensionality reduction and clustering techniques were a similar set of analyses were employed as before with the 2D data for comparative statistics among samples.

*Visualization*–Data visualizations, including PCA and UMAP scatter plots with embedded thumbnails of object annotations, were generated in a similar manner as in the 2D measurements.

*Data saving and management*.—Measurement metrics were saved in multiple formats to accommodate various analysis needs. Individual metrics were stored in text files, while aggregated data was exported to CSV and JSONL (JSON Lines) formats using Python’s *csv* and *json* libraries, respectively.

### Color and Color Pattern Extraction

*Color segmentation and hue normalization.*—In order to, enhance and normalize the color of segmented biological images a python script needed to be developed in order to make the extraction of color and color pattern more accessible. This approach involves four main steps: color segmentation, hue normalization, color extraction, and optionally color pattern alignment. Below, we detail the steps and functions used, referencing the relevant tools and techniques.

The *recolorize* (Weller et al., 2022) package served as inspiration in part but the author needed to develop a python implementation for many similar functions in order to more easily bridge python computer vision and machine learning with color analyses of biological specimens. While similar to the *recolorize* package in *R* (R Core Team, 2021; Weller et al., 2022), the work here is from the ground up a novel *python* recipe for hew normalization and color segmentation using a variety of pre-existing functions.

*Reading Image and Mask:* The input image and the corresponding binary mask are registered using the *skimage* library (Van der Walt et al., 2014). The binary mask is applied to the foreground mask image from the previous step to isolate the region of interest. Pixels outside the mask are set to zero. The RGB colors can then be captured from inside the mask using it as an RoI to produce a fully color RoI.

*Felzenszwalb and Huttenlocher segmentation (FHS) of color.* —The ‘Color Segmentation’ button applies FHS to the masked image to identify different regions based on color and texture (Felzenszwalb & Huttenlocher, 2004). This algorithm is widely used due to its efficiency and ability to produce meaningful segmentations that preserve object boundaries. The algorithm treats the image as a graph, where each pixel is a node, and edges between nodes represent the similarity between adjacent pixels (Felzenszwalb & Huttenlocher, 2004). The goal is to partition this graph into segments that correspond to distinct regions within the image, in this case colors. The FHS segmented image is modified based on the average color of each segment using *felzenwalb* from the *skimage.segmentation* and *skimage.color.label2rgb* libraries (Van der Walt et al., 2014). The FHS image is saved to the specified path using *skimage.io.imsave*.

*Color normalization.*—To normalize color for downstream color analyses and enhance the visual quality of images a custom function *normalize_color* was written needed be written. The input image is read using *cv2.imread* and converted from BGR to LAB color space using *OpenCV* (OpenCV Contributors, 2021). The LAB color space, also known as CIELAB, is designed to be perceptually uniform, meaning that a given amount of numerical change corresponds to roughly the same amount of perceived color change(Sharma & Trussell, 1997; OpenCV Contributors, 2021). The LAB color space is widely used in various image processing tasks, including color correction, image segmentation, and texture analysis (McLAREN, 1976; Sharma & Trussell, 1997; OpenCV Contributors, 2021). LAB stands for, L (Lightness): represents the lightness of the color, ranging from 0 (black) to 100 (white); A (Green-Red): represents the position between green and red colors, with negative values indicating green and positive values indicating red; B (Blue-Yellow): represents the position between blue and yellow colors, with negative values indicating blue and positive values indicating yellow (McLAREN, 1976). The LAB color space is particularly useful in image processing because it separates the lightness component (L) from the color components (A and B), making it easier to manipulate color and brightness independently. This separation allows for more precise adjustments and is particularly beneficial in tasks such as color normalization and contrast enhancement (McLAREN, 1976; Sharma & Trussell, 1997).

Contrast Limited Adaptive Histogram Equalization (CLAHE) is applied to the L channel to enhance the contrast of the image (Zuiderveld, 1994; Bradski, 2000). CLAHE is an improved version of Adaptive Histogram Equalization (AHE) that aims to prevent the over-amplification of noise in homogeneous regions of the image (Pizer et al., 1987; Zuiderveld, 1994). CLAHE works by clipping the histogram at a predefined value (the clip limit) before applying the histogram equalization. This prevents the over-amplification of noise and ensures a more balanced contrast enhancement across the entire image (Pizer et al., 1987; Zuiderveld, 1994; OpenCV Contributors, 2021). The image is then converted to BGR encoding and later to RGB encoding for visualization. The normalized image is saved to the specified path using cv2.imwrite.

The FHS or hue normalization can be run together or independently. This methodology employs advanced image processing techniques to enhance the color and texture features of insect images, facilitating better visualization and analysis.

*Average color vector analysis.*—Each RoI was reduced to a single average color vector in the LAB color space. This method involved flattening the LAB values of the entire RoI and computing the mean across all pixels, resulting in a 3-dimensional vector representing the average L, A, and B values for each specimen. Although this approach is computationally efficient, it has the significant drawback of losing all spatial information about the color distribution and is not invariant to differences in shape.

*Shape invariant color pattern analyses.*—To perform shape invariant pattern analyses constructing a 3D color histogram, or "color cube," in the LAB color space for each RoI was employed (Bradski, 2000; OpenCV Contributors, 2021). Each RoI, was isolated using binary masks to exclude any background pixels. For each RoI, a 3D histogram was calculated that quantified the distribution of colors across the L, A, and B channels . The histogram was divided into bins, with each bin representing a specific range of colors within the LAB space. This histogram effectively captures the frequency of occurrence of each color within the RoI, independent of its spatial location. By focusing on the distribution of colors rather than their arrangement, the color cube method is inherently invariant to the shape of the sclerite or category in question. To facilitate subsequent analysis, the 3D histogram was flattened into a 1D vector, representing the overall color distribution within the RoI. This step transforms the complex color patterns into a standardized format, enabling direct quantitative comparisons between specimens.

*Principal component analysis (PCA) on color histograms.*—Following the generation of color histograms, PCA was employed to reduce the dimensionality of the histogram vectors and to identify the principal components that capture the most significant variation in color distribution across the specimens. Prior to PCA, the histogram vectors were standardized using Z-score normalization to ensure that each feature (i.e., each bin of the histogram) contributed equally to the analysis.

The PCA was performed on the normalized histogram vectors, reducing the multidimensional color data to two principal components (PCs). These PCs were then plotted to visualize the clustering of specimens based on their color patterns. Importantly, since the input to PCA is derived from the color histograms—which do not encode any shape information—the analysis remains focused solely on color distribution, free from any influence of shape.

*Extracting color information from the RoI.*—Using *OpenCV, sklearn*, and *numpy* specialized function was developed calculate the average color of the masked region in the image (Bradski, 2000; Pedregosa et al., 2011; Harris et al., 2020a; OpenCV Contributors, 2021). To do so the binary mask was applied to the original image, extracting the pixel values where the binary mask is white (mask == 255), and then computing the mean color of these pixels.

Sometimes it is useful to just know the most frequent color in a region. Via another new function, the *dominant_color* function, written to find the most frequent (dominant) color in the masked region. It applies the mask, reshapes the masked image to a list of pixels, counts the frequency of each color, and returns the most common color. Using KMeans clustering via *sklearn* (Pedregosa et al., 2011) to find the top k colors in the masked region another function applies the mask, reshapes the masked image, performs KMeans clustering, and returns the cluster centers (top k colors) and the labels for each pixel. The RGB values for the most frequent color, average color and the top k colors are saved to an output file. These RGB values can then be used downstream for quantitative analyses of color.

*Converting extracted colors to human-readable names.*—A color lookup table maps RGB color values to human-readable color names. This table includes a wide range of colors, particularly focusing on browns and tans. The RGB colors in the lookup table are then converted to the LAB color space. LAB color space is used because it is perceptually uniform, meaning small changes in LAB values correspond to consistent changes in perceived color. A KDTree was built using the LAB color values via *scipy spatial* library (Virtanen et al., 2020). The KDTree allows for efficient nearest neighbor searches, which are used to find the closest color in the lookup table to any given LAB color. A custom function*, closest_color_name*, converts an RGB color to LAB color space, make queries to the KDTree, find the nearest LAB color in the lookup table, and returns the corresponding human-readable color name or the nearest value to name match in the lookup table. The results are written to a new text file, including the human-readable color names. Finally, to incorporate ranges of both the human readable colors and the color histograms per group an optional grouping file can be used to provide a method of concatenation by group.

### Integration of Instance Segmentation and Morphometrics with VLMs to Produce Species Descriptions

*Adding expert morphological information to images for GPT-4V.*—To pass the information already gathered from the instance segmentation, morphometric and color segmentation scripts as well as taxonomist defined questions needed to produce a species descriptions *Descriptron* contains functionality to help users prepare data and perform batch VQA tasks with *GPT-4*.

We decided to overlay the contours from the instance segmentation masks over the images to help direct the *GPT-4V*. With *Descriptron*’s ‘Visualize Contours & Bbox’ button the user can also decide which category (eg. tibia, wing etc.) of instance contours they wish to overlay on the images, or they can overlay all of them with a different color for each as an option. The category name will also be displayed in the same color as the contour. Additionally, the user can define the opacity of the contour as well as the color. This can allow the user potentially to make composite images of different contours that may overlap to a great degree with one another. This is important as it will allow the user a lot of downstream control on how they wish to present morphological features to the VLM for analyses.

*Prompt generation specifically for fine tuning GPT-4V.*—In order to handle a single to hundreds or thousands of specimens we developed a script that converts user generated prompts about the important morphological features in user images. This is where previous taxonomic knowledge is required in order to request back the required information from the image needed to complete a species description. We give the user the option to fine-tune *GPT-4V* using a simple tab separated file, containing the image path, question and answer. This is converted into a JSONL via the ‘TSV to JSONL’ button for fine-tuning *GPT-4V*. To merge the image file (png or jpg) the script opens the image file in binary mode and the *base64* library converts binary to base64 and then decoded to UTF-8 for embedding in the JSONL file (Foundation, 2023). This step allows the users to eliminate the need for external reference files. This step also sets the initial prompt scenario for the VQA task and then adds in the image and the prompt questions.

*GPT-4V fine tuning and GPT4.*—We use the *openai* python library to load the JSONL file for commands to for the commands from our JSONL file to the OpenAI fine tuning process (OpenAI, 2023a). OpenAI recommends approximately 50–100 well-crafted examples for fine tuning but the final sample size depends on the model of choice and task (OpenAI, 2023b; Open AI, 2024). *Descriptron*’s ‘Fine-tune GPT-4o Model’ directly interacts with OpenAI to pass the JSONL file through OpeanAI’s fine-tuning process.

The ‘GPT-4 Featurize’ button will then interact with either the *GPT-4V* model, other GPT models or our fine-tuned model from the previous step can also be used. The available models at the time of writing this manuscript with *GPT-4V* for fine-tuning are gpt-4o (gpt-4o-2024-08-06), and gpt-4o-mini (gpt-4o-mini-2024-07-18), and for image inference gpt-4o, gpt-4o-mini, or gpt- 4-turbo (Open AI, 2024). The ‘GPT-4 Featurize’ button converts a user provided tab separated values file of questions and converts them into a JSONL format. Additionally, it the *base64* module converts the specified input images into base64 format so that the files can be sent to OpenAI and processed using the users OpenAI API key. The model’s response is stored as a plain text file or optionally as a JSON file, for downstream processing. If a folder of images is provided the script iterates through all image files applying the workflow to each image. With the ‘GPT-Featurize’ button the user can provide guided questions about biological images to obtain descriptions of large datasets of biological specimens using standardized queries.

*Automating materials examined section*.—Specimen labels with clear writing can have joint OCR and NLP tasks performed via *GPT-4V* which performs well at these VQA tasks (Shi et al., 2023). The ‘GPT-4 Specimens Examined’ loads user defined images of specimen labels along with question prompts about the label. The example question prompt contains the Darwin core headers as part of the question so that response data can be parsed downstream. If working from type material, there is an option to input that directly if the type label is present this will also get processed by *GPT-4V* and added to the output if present. If no type status is indicated the script defaults to ‘Specimens Examined’. Results are saved in JSON format for downstream processing. The ‘Parse Material Examined’ button extracts the critical data from the previous step and concatenates it, into a coherent description of the locality removing the Darwin Core headers so that it reads more like a typical species description.

*Concatenation of morphology characters and species description refinement*.—This step brings together the various data sources we have been collecting so far on our specimens. Using the ‘*Concat Description*’ button *Descriptron* runs a script that takes advantage of the structured file names that we have been producing at each stage as output we can link categories to specific species using the base file name from a user specified list of species or samples to concatenate. The different morphological features from the *GPT-4V* section, the color data, and morphometric data, and the material examined data are then concatenated. All this data is gathered for downstream processing so that we can prompt in the morphological data of each sample along with the measurement ranges by species or specimen group.

*Generating an initial species description*.—With the ‘*Generate Species Descriptions*’ button we prompt in the concatenated sample or species data from the previous step to OpenAI using their *openai* python module and our API key (OpenAI, 2023a). As we are only prompting text this time we can take advantage of other models with superior reasoning capabilities such as o1-preview and o1-mini (OpenAI, 2024a). Using these simple plain text examples of questions and answers for the LLM, users can prompt with specific examples of how they want the output to look in terms of format as well as specific requirements for the ‘Diagnosis’ section. The diagnosis section can now be prompted in and the LLM model can try to perform inference about a logical response for the user’s requirements, for a given ‘Diagnosis’ section. We provide a plain text file example that users can modify for their own prompts which are then passed to OpenAI via an *argparse* statement or via the dialog box window in the GUI. As the species descriptions can be very long and above the default token length the script automatically parses the input file into chunks that are below the token length of 8,000 tokens so the files can be processed. We also take care never to cut a species description so that we do not need to implement a sliding window approach. Each chunk is read in before requesting the final response.

*Generating a dichotomous key*.—Using the output from the previous step we take a similar approach to read in the input in chunks. Users can provide input as a simple text file of a dichotomous key from a similar taxonomic group or for a specific output format to be followed. The prompt which includes the user provided examples is then passed to OpenAI where the user specified *GPT* model performs inference, structured output can also be requested. *GPT-4* has been trained and performs well at delivering structured output tasks and has been trained on both structured and unstructured data (OpenAI, 2024b). In this initial version we do not activate the structured output method but future versions we plan to implement this approach and compare to unstructured output.

*Species description evaluation methods*.—To compare species descriptions generated by *GPT-4V* or by various VLMs and or prompting recipes *Descriptron* has a series of analyses that can be performed on the species descriptions themselves. ROUGE (Recall-Oriented Understudy for Gisting Evaluation) is a set of metrics commonly used to evaluate the quality of machine-generated text, such as summaries, against human-written references (Lin, 2004; Pltrdy, 2017). It primarily measures overlap between n-grams, word sequences, or word pairs between the generated text and reference text (Lin, 2004). The key variants include ROUGE-N, which calculates the overlap of n-grams, and the ROUGE-L, which uses the longest common subsequence (LCS) to evaluate fluency and structure (Lin, 2004). A higher ROUGE score indicates better alignment with the reference text, making it a valuable metric for assessing both the informativeness and precision of generated outputs (Lin, 2004). For visualization we provide synteny graph network plots of the most common adjectives and nouns with their most common connected nouns or adjectives. Common noun and adjective histograms and Venn diagrams are also provided by *Descriptron* as part of its species description analyses buttons.

Another way *Descriptron* provides a visualization of the similarity between species descriptions is through dimensionality reduction of the VLM embeddings. PaCMAP (Pairwise Controlled Manifold Approximation) is a dimensionality reduction method that can be used for visualization, preserving both local and global structure of the data in original space (Wang et al., 2021). For VLMs, embeddings are crucial for aligning information from both modalities (vision and language) in a shared embedding space. Unlike other dimensionality reduction techniques like t-SNE and UMAP, which focus predominantly on either local or global neighborhood structures, PaCMAP balances these two aspects (Wang et al., 2021). This balance allows PaCMAP to represent (1) Multimodal Representations: VLMs, such as CLIP or GPT-4V, generate embeddings for text and images that can be directly compared, enabling tasks like image-text retrieval, captioning, or multimodal reasoning; (2) Alignment in a Shared Space: Embeddings encode semantic meaning, ensuring that visually similar images or text with similar semantic content are positioned closer in the embedding space (Wang et al., 2021; Huang et al., 2022). For example, a photo of a dog and the word "dog" might have closely aligned embeddings. This second aspect makes it useful for plotting the joint embeddings from both images and the description. To accomplish this *Descriptron* takes the image used for the species description section along with the description and extracts the embeddings from the encoding from CLIP (Radford et al., 2021). We implement a sliding-window approach to generate embeddings for the species descriptions as they are significantly longer in general than the token limit of CLIP (Radford et al., 2021). Using the embeddings from the image and text of the species description we provide a quantitative method to evaluate taxonomic species descriptions and their accompanying images together for the first time. In addition to the PaCMAP analyses we also perform a cosine similarity measure on the embeddings. The cosine similarity measure in our implementation is the joint vector of the image and text (from the Description section). The cosine similarity measure scores how similar the two descriptions are in vector space. Cosine similarity measures how the vector of the embedding overall point in the same direction by normalizing the magnitude (Li, Koh & Du, 2024; Steck, Ekanadham & Kallus, 2024). This allows us another method to quantify the similarity of the two descriptions (e.g. Taxonomist vs GPT-4o) and compare species descriptions by species as well as by overall difference between the two treatments.

### Initial Testing of Descriptron Via Describing Psylloidea Forewings

*Study system.*—Psyllids or jumping plant lice (Hemiptera: Sternorrhyncha: Psylloidea), are plant-sucking insects, with several species being economically important pests, which transmit pathogenic bacteria and cause serious plant diseases. This superfamily comprises approximately 4,000 described species, with about the same number of undescribed species, primarily inhabiting temperate and southern temperate regions. Despite their economic importance, psyllids remain understudied due to their small size and taxonomic complexity. The model groups of the current study are the psyllid genera *Melanastera* and *Russelliana*, which belong to the two different families, Liviidae and Psyllidae, respectively. Numerous new species from these genera were recently described from the Neotropics (Serbina & Burckhardt, 2017; Serbina et al., 2024).

*Model preparation and annotation using Descriptron.*—One of the key morphological features needed to describe new species of psyllids is the shape of their forewings and the pattern of their wing venation. Many psyllids contain wing venation that are nearly obscured by coloration pigments. This makes it very difficult to use methods that rely on contrast, even novel advances in recent methods that rely on advanced computer vision alone might struggle here (Eshghi et al., 2022). Models that rely on subtle differences that can be learned could perhaps make this task possible in an automated manner. We used *Descriptron* to annotate ten different wing cells along with the entire forewings in the genus *Melanastera*. Briefly we used 40 different species of *Melanastera* (including males and females from several species to include sexual dimorphism in the training data) utilizing one individual right forewing from each species. A complete list of characters can be found in Appendix 1. Using a combination of *Descriptron*’s implementation of *SAM2* (Ravi et al., 2024) and hand-crafted contours using *Descriptron*’s ‘Keypoints to Mask’ button and the eraser and paintbrush tools. To examine the utility of the *Detectron2* (Wu et al., 2019) we used *Descriptron*’s default settings to run *Detectonr2* but with the maximum number of iterations set to 10,000 and evaluation set to every 1,000 iterations.

Next, we performed inference on 48 individuals from 43 species of *Russelliana* including both males and females from five species to include sexual dimorphism in the test data. Many of the wing veins in psyllids are similar; however, the pterostigma (an enlargement of the costal wing vein) can be greatly reduced in some taxonomic groups. The *Detectron2* model performed well on many of the cells but often either did not predict the ‘pterostigma’ correctly or left it out entirely. The authors adjusted the predictions to include the pterostigma and other errors using the *Descriptron* paintbrush and eraser tools as well as the ‘Keypoints to Mask’ and the ‘Re-annotate Mask’ buttons.

*Morphometric, geometric morphometric, and color analyses*.—To include both measurement data and to capture color data in human readable format we utilized *Descriptron* to capture the width and length of the different forewing cells. In addition to this we also collected the geometric morphometric data of our wing cells and the forewing. Finally for each forewing category we also collected the color as an average value and the most dominant color of each category as well as quantifying the color pattern as a color histogram using the default settings of *Descriptron*.

*Concatenation and species description generation*.—The instance segmentation masks from the *Descriptron* were visualized using the ‘Visualize Contours and Bbox’ button. The label data was also simulated using a locality label for one of the species to mimic a batch collection and so that label data would remain the same using the ‘GPT-4 Specimens Examined’ and ‘Parse Materials Examined’ buttons. The species descriptions section were generated using custom prompts that first used a ‘seed prompt’ setting up the scenario for the agent along with two examples of the structure and type of output the user required, examples available from https://github.com/alexrvandam/Descriptron (Van Dam, 2024). In addition to the seed prompt there were 24 individual questions needed to complete a thorough species description of the psyllid forewings using the ‘GPT-4 Featurize’ button. Next the individual questions were refined further, and a ‘Diagnosis’ section was written using the ‘Generate Species Descriptions’ button using the default *GPT-4o* model. Following this a dichotomous key could be written using the ‘Refine Output and Dichotomous Key’ button. All prompts can be found in the *Descriptron* GitHub page (https://github.com/alexrvandam/Descriptron) (Van Dam, 2024). Finally, the similarity in the species descriptions were manually evaluated for errors and taxonomist generated descriptions were compared to those of *GPT-4o* using *Descriptron*’s ‘Calculate Rogue Stats’ button. A subset of the categories were also not included in the species descriptions and we wanted compare the effect of not including the contours and labeling of wing cells to those that had all the characteristics outlined and labeled in the image. To do this a Shapiro-Wilk test was performed followed by a Wilcoxon Signed-Rank using *scipy* test to identify if labeled vs unlabeled categories had more errors (Virtanen et al., 2020).

*AI use statement.*— AVD used GPT-4o (OpenAI et al., 2024) to assist in troubleshooting and generating initial code from time to time, the author found it extremely useful.

## Results

### Qualitative Evaluation of Descriptron’s Morphometric, Geometric Morphometric Color Measurement Tools

*Geometric morphometric clustering.*—The contour extraction was successful and it can be queried for specific sclerites and families, even down to specific individual specimens. The ordering of the points and converting the contours to semi-landmarks was effective and the distribution of points can be seen to close large gaps in the initial contours compared to semi-landmark conversion. Initial principal component followed by Procrustes transformation was also successfully performed by *Descriptron*. Geometric morphometric clustering performed as expected and was able to perform semi-landmarking on wing cell categories. This allowed for analyses of the overall shape of the entire forewing as well as for each individual wing cells. Please see Figures 1 and 2 for an example output the statistical clustering output.

*2D morphometric measurements.*—A few of the scale bars were not identified by the method utilizing *CV2* (Bradski, 2000). This happened when there were also straight and long wing veins near the scale bar and since the method implemented in *Descriptron* looks for scale bars close to the contour sometimes it picked up on the contour itself. When the wings were more rounded or had significant pigmentation the scale bar was successfully identified. To make the measurement uniform we decided to just use the scale in pixels to be added in manually into the *Descriptron* dialog box as many of the images had the same scale to pixel ratio.

*Color extraction.*—The color analyses performed as expected and we were able to extract the color for each cell and for the entire forewings. The color histogram analyses also performed as expected, and we were able to extract metrics for each category. The color histograms were then also plotted as a PCA and the script performed as expected.

### Quantitative Evaluation of Descriptron’s Species Description Tools

*Quantitative similarity of word choice and sentence structure of GPT-4o and taxonomist species descriptions.*— Nine of the specimens did not fully complete the bioinformatic pipeline probably due to a text file parsing error, thus the statistical analyses are based on 43 descriptions in total. Despite trying to constrain the model to a specific text format *GPT-4o* often formatted different sections slightly making it difficult to capture all its output variation through uniform hardcoded data carpentry in *python*. Future versions of *Descriptron* will utilize *GPT-4o* structured output mode using JSON format so that text based parsing errors do not occur. The average and standard error for the ROUGE-1, -2, and -L were 0.45±0.01, 0.11±0.00, and 0.22±0.00 respectively (Supplemental Figure S1, S2). In terms of total adjective and noun word choice across both methods the taxonomist and *GPT-4o* descriptions were 26% and 27% similar (Supplemental Figure S3). The taxonomist species description shared 37% and 52% of adjectives and nouns with the *GPT-4o* species description respectively (Supplemental Figure S3). In terms of the adjective and nouns used for the *GPT-4o* description it shared 51% and 39% with the taxonomist description respectively (Supplemental Figure S3). A full breakdown of all the ROUGE scores is summarized in Supplemental Figure S2 and the most common adjectives and nouns can be found in Supplemental Figures S4 and S5 respectively.

In addition to ROUGE scores, we also generated synteny plots of the word choice (Fig. 3). The 50 most common adjective-noun pairs and in another synteny plot the unique adjective-noun pairs. The nodes are either nouns or adjectives and the edges represent the co-occurrence in the same sentence (Fig. 3). The weight (represented by color) is the frequency of co-occurrence and nodes are occurrence (Fig. 3). There seemed to be differences in word choice between both the top-50 and unique words, however these differences did not produce a statistically different graph of the combined data with a Wilcoxon density, average clustering and average degree of 0.0 with a P-Value of 𝑝=1.0, indicating the overall structure of the synteny graphs were constructed was not statistically different.

**FIGURE 3.**
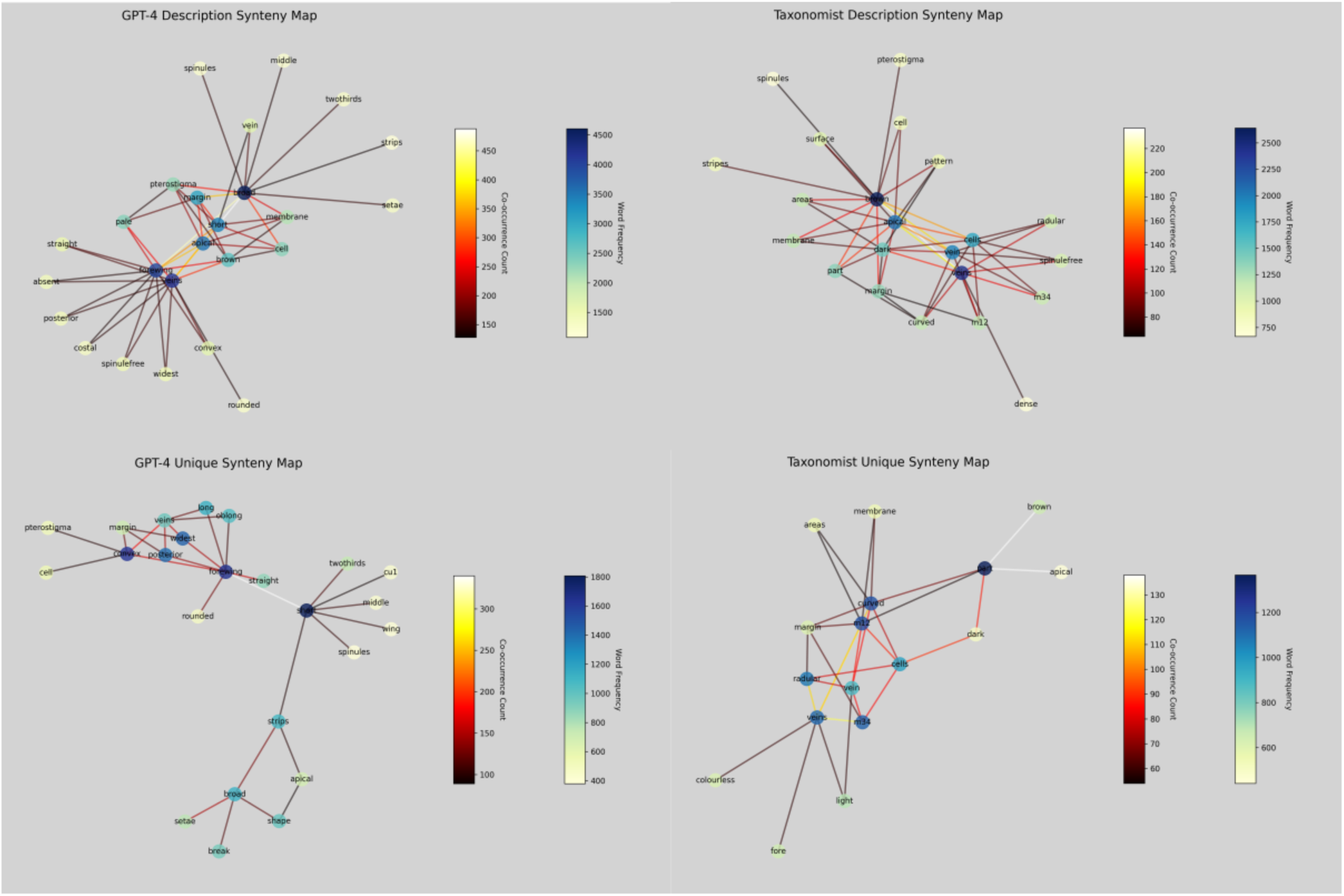
Left column *GPT-4o* fine-tuned model species descriptions, right column Taxonomist produced species descriptions with top row being synteny maps of common word pairs and bottom row unique word pairs synteny maps. Species descriptions were produced from 43 *Russelliana* species descriptions produced from outputs and prompts and passed to *GPT-4o* OpenAI API using *Descriptron*.

*Overall similarity of joint image and species descriptions by GPT-4o and taxonomist.*—The average cosine similarity and standard error was 0.94±0.00 with a range from 0.90–0.96 (Supplemental Figure S6). The PacMAP analyses (Figure 4) however shows that the two groups of species descriptions are quite distinct clusters indicating that the embeddings from the descriptions produced by the two methods were semantically different species descriptions.

**FIGURE 4.**
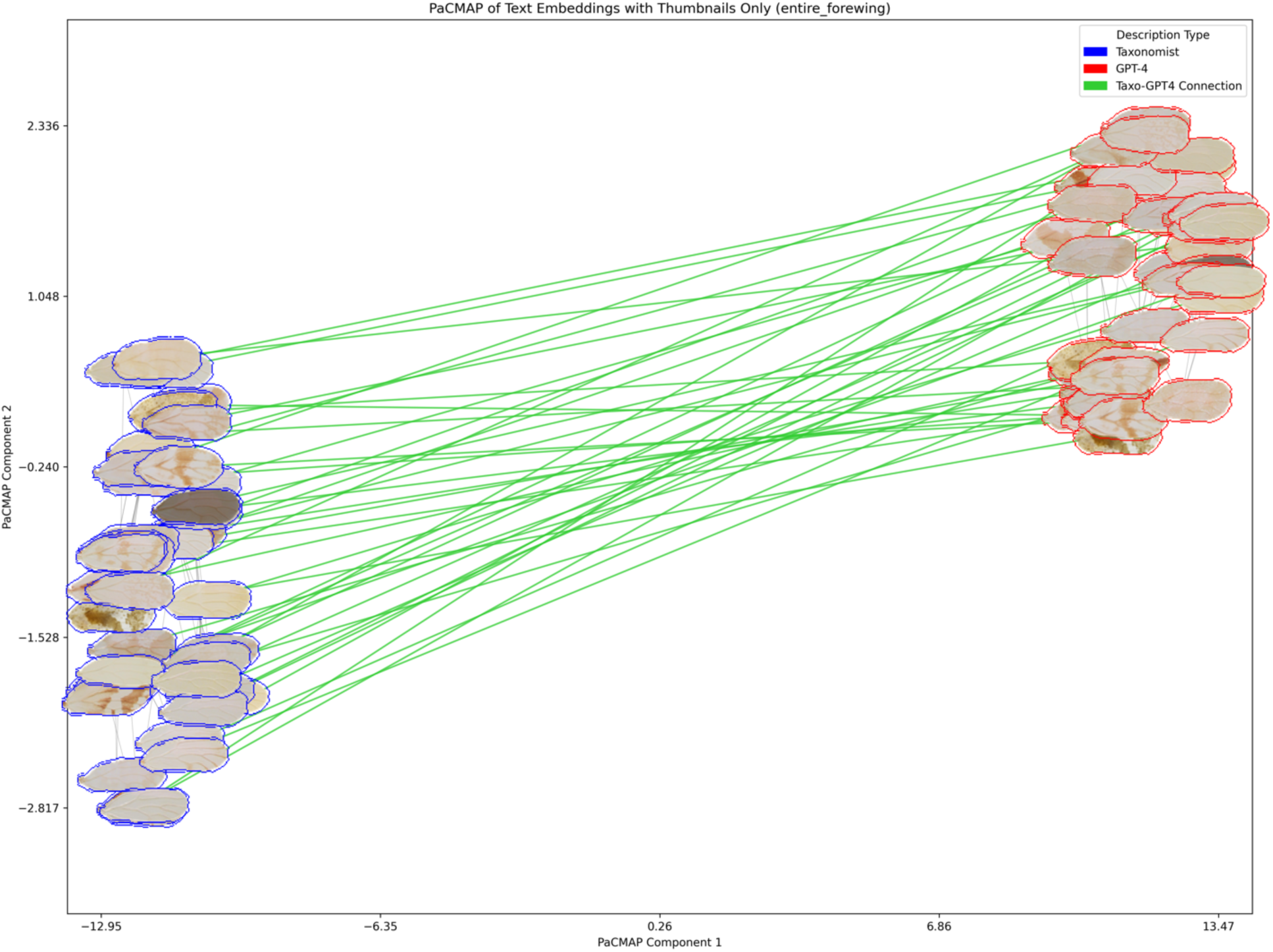
Quantification of taxonomic species descriptions using joint image and text embeddings, plotted via PaCMAP clustering. The embeddings are based on annotated images and ’Description’ sections from taxonomic species descriptions of *Russelliana* forewings, produced by expert taxonomists and *Descriptron*, a fine-tuned *GPT-4* model. Each forewing is positioned in the PaCMAP plot according to its principal component coordinates. Long green lines connect embeddings of the same species, while thin gray lines indicate the three nearest neighbor species embeddings.

*Diagnosis and dichotomous key*– As the keys were generated prompting the use of the ‘Diagnosis’ sections they should approximate the valuable characters found there. The taxonomist generated key contained many more characters as the original analyses was not restricted to the forewing (Serbina & Burckhardt, 2017) making direct comparison difficult. However, the overall structure of the dichotomous key was consistent and the use of characters that the model perceived to be present seemed logical.

*Evaluation of similarity and errors in Descriptron generated species descriptions to expert taxonomist descriptions.*—*GPT-4o* made on average 5.52±0.20 (average±standard error) mistakes or hallucinations per species description across the 43 specimens examined. Individual errors in the descriptions themselves were often due to not including instance segmentation of a given feature. Character annotation vs characters lacking annotation was first examined to see if the errors were normally distributed by a Shapiro-Wilk test for normality and it rejected that both sets were normally distributed. As the data was not normally distributed a Wilcoxon Signed-Rank Test was used to see if there was a difference in the number of errors that had labels vs those that did not. The Wilcoxon Singed-Rank test yielded a W=2.35 and a P-Value: 𝑝=0.05 indicating that there was only a weak difference between the characters that were had contour labels and those that did not. In many cases the description left out the ‘pterostigma’. This character often had the label obscured by the segmentation line which probably made it difficult for *GPT-4V* to accurately read the label, effectively making this an unlabeled category.

## Discussion

### Future Improvements and Caveats

Both the ROUGE scores and PaCMAP clustering indicate that the species descriptions produced by the *Descriptron* were distinctive compared to those that a taxonomist would write (Figs. 3,4). The descriptions produced by *Descriptron* also contained a few key errors for each species requiring user correction. This indicates that while automated this first attempt still requires user supervision to produce adequate species descriptions. Despite this needed supervision and error correction much of the species’ descriptions were still quite accurate. The tradeoff between the good quality data produced by *Descriptron* and the time needed to correct its errors was a significant net benefit in timed saved for this task. In about an afternoon worth of editing most of the errors could be fixed for 43 species descriptions. This is a significant amount of time saved compared to handwriting them from scratch. The number errors were weakly significant indicating that perhaps inclusion of annotation of additional characters on images or structures would have further improved results. There is some evidence for this from other studies looking to optimize VLM performance (Guo et al., 2024; Wan et al., 2024). One possible solution to providing additional information would be to present composite images of the same image with a few of the key morphological features segmented in each subframe. In this way many more morphological features could be given to the VLM without obscuring the underlying specimen image or category labels.

Fine tuning could also help improve the results. Here we develop the capability for the user to fine tune *GPT-4o* with vision on both text and images. In the present study we looked at the standard *GPT-4o* model without fine tuning. Future work should experiment with both the number of fine-tuned examples, and if labeling images with contours and text helps improve the VLM’s understanding of the subject matter further. Optimizing VLM’s for VQA tasks is an active area of research in the AI field currently and taxonomy has something to contribute to the advancement of this field. *GPT-4o* is one of the frontier models in AI currently and scores quite high on math and science recall (OpenAI et al., 2024; Stribling et al., 2024). However, with answering taxonomic questions which are primarily visual it seemed to struggle and be pushed to its limits on what text it could reliably return without it being rife with errors. This is why we highly recommend checking its responses, they should be thought of as a first rough draft that will require significant editing for taxonomic purposes. Future work should also plan to provide a standard benchmark taxonomic dataset to help evaluate alternative VLMs to compare to GPT- 4o.

Including more data from the measurements and color extraction as part of the initial prompts for the species description could also help reduce errors. *Descriptron* should provide a reliable platform for further testing and optimization. Additionally other VLMs could be tested and compared as well for this task. We did not include measurement images or data in the initial prompts, in the final prompts for refining the output and generating the dichotomous keys we did, but it appears to have been too late at this stage and simply as much data as possible should be given in the initial prompts.

The 3D measurement tool needs further testing and improvement before it can be more reliable, but we release it here so that this process can begin. Probably the best way to use it is by providing a user generated depth map from stacked images that included depth data. Some microscopy systems provide such depth maps with their stacked images. Future work should focus on allowing users with other focus stacking setups that utilize standard macro-photography cameras a way to provide accurate depth maps. In addition to this stereo imaging through the rotation of the camera position around the specimen or through novel view synthesis using multiple views might also provide more accurate 3D depth and there by 3D measurements.

*Descriptron* provides many automated tools for taxonomists or scientists of any background (both amateur and professional) to collect data on their taxa of interest and test evolutionary hypotheses. The instance segmentation workflows using *SAM2*, *Detectron2*, or simply the *Descriptron* paintbrush tool, allowing users to rapidly record instance segmentation of regions of interest which can be communicated to a VLM through image annotation. The morphometrics workflows allows users to gather measurement data from a given region of interest. Morphometric data such as size and relative dimensions are commonly used in evolutionary biology hypothesis testing and in taxonomy. The geometric morphometric workflows also allow users to cluster specific regions of interest to identify if specific taxa contain significantly different shapes. Again, differences in shape are frequently interrogated in both taxonomy and more broadly in evolutionary biology. Color can also be quantified using the tools in *Descriptron*. Differences in color can be interrogated via clustering using *Descriptron* or can be directly exported as and utilized in other evolutionary studies as categorical or continuous data. The 3D morphometrics for now is in its experimental stages and needs validation, but we expect it to become both reliable and useful as the other workflows with more thorough validation. All these tools can be used to help elucidate differences between taxa. These data once exported can also be pasted into prompt templates and used in the semantic species description workflow phase using a VLM. Species description workflows with *Descriptron* allow the users to modify prompt templates and experiment with different ways to prompt in data to *GPT-4o* using the OpenAI API. This allows for relatively cheap experimentation with different image annotations and prompts to find the ideal method for describing their focal taxa in a repeatable and automated manner. Then species descriptions can be compared to identify if different input prompting practices or instance segmentation approaches produce significantly different results. This will allow users to dial in on improving the taxonomic validity of the species description output for their group. A very similar program TaxonGPT takes input form of Nexus matrices, which contain species and their corresponding character states (Huang et al., 2024). TaxonGPT uses text files of characters and character states to help constrain GPT-4 from generating inaccurate information by using knowledge graph semantic representations (Huang et al., 2024). This integration ensures that output from TaxonGPT generates taxonomic descriptions and keys that are grounded in accurate and structured data (Huang et al., 2024). This seems to be a very promising approach. A similar approach could be taken by *Descriptron* in the future to check for any errors by feeding in hard coded data from the instance segmentation as an error checking process.

We hope to add even more capabilities to *Descriptron* in the coming year including keypoint prediction and associated geometric morphometric techniques that use keypoints. We also plan to improve the user friendliness and provide updated workflow ‘how to’ instructional documents and videos. Additionally, we plan to expand the 3D capabilities to include methods such as stereo vision and novel scene prediction. The file size of ‘*Descriptron*’ currently is also quite massive at around 137GB as we include all the models and data sets needed to prevent version incompatibility issues. We plan to continue to reduce this size of *Descriptron* and different ways to download the *Descriptron* in the future for advanced users. However quite a few modern video games are a similar size and this has not stopped their popularity (Park, Morgan, 2023).

### Descriptron Provides a Significant Time Savings for Taxonomic and Morphological Work and Quantifies Taxonomic Species Descriptions

This is the first study to attempt to automate species descriptions using VLMs and other deep learning tools. Despite the many improvements that can be made *Descriptron* is a huge time saver for taxonomists. The time needed to produce similar species description data would have taken multiple weeks to months but was generated in less than ten days including training the CNN for Detectron2 and editing of *GPT-4o* output. We hope this tool can be used widely by both amateur and professionals alike to speed up biodiscovery and the description of new species.

VLM’s are the current state of the art in both encoding and decoding of the important regions of a given image into text. Despite this, the level of detail needed to describe species requires expert taxonomic knowledge to guide these models. The results from the present study seem to indicate that expert taxonomic knowledge is needed to help inform these models placing taxonomists (again both amateur and professional) in a well-situated position to harness these new tools to speed up the formal description of novel species. *Descriptron* will help in this endeavor by providing a reliable GUI to allow taxonomists who may not have extensive experience in computer coding a way to form new methods and hypotheses about how to use VLMs to interrogate their datasets of putatively novel species. By providing taxonomists a new tool to leverage their knowledge, *Descriptron* enhances their ability to explore and describe biodiversity, supporting their expertise in the process.

Here for the first time, we also offer a way to quantify both the images and text of key morphological features found in taxonomic species descriptions either one at a time or jointly by using the VLM embeddings and associated statistics (eg. cosine similarity scores, PaCMAP coordinates). The ability to quantify a whole variety of new colors, textures and shapes of important morphological characters will in turn allow for new methods of analyzing them using continuous variables in statistical inference, besides being able to judge the similarity and repeatability of taxonomic species descriptions in a statistical framework.

The significant upfront time in developing models to extract morphological features if focused on just the top 20 most diverse clades could be a way to significantly improve our knowledge of ‘Dark Taxa’ and there by a significant proportion of the Earth’s biodiversity (Srivathsan et al., 2023; Meier et al., 2024). That same upfront time needed to train specific instance segmentation models for key morphological traits might not be worth the time if clade diversity is low and the time needed to error correct *Descriptron*’s output is too costly. We suggest based on the initial findings in this study that *Descriptron* could be used to build up a model suite capable of delimiting key morphological features to help constrain VLM’s with annotated images. This in turn would offer a huge time savings for the most species rich clades of ‘Dark Taxa’. Obviously more improvements as to how the image annotations could be presented to the VLM will improve the draft species descriptions, but that is what makes this a new and exciting avenue of research with a large potential payoff in terms of being able to produce species descriptions in a rapid automated way.

We hope *Descriptron* will be used by taxonomists and evolutionary biologists alike to describe new species and collect new morphological data. This in turn provides an improved understanding of our planet’s biodiversity deepening our understanding of the complexity and interconnectedness of life on Earth.

## Acknowledgements

AVD would like to thank his family (especially Arnav who was very encouraging about this project from the beginning) for allowing him to move everyone to Germany to complete this project. AVD would like to thank Dr. Sean Locke (University of Puerto Rico Mayagüez) for initial discussions that helped inspire this project and grant proposal proof reading and for helping to think up the name *Descriptron*. We thank Dr. Rudolf Meier for initial review of the manuscript and for developing positions in Taxonomy at the Museum für Naturkunde Berlin (MfN) specifically to work on dark taxa biodiscovery and automated descriptions without the support of MfN the completion of this project would not have been possible. We thank Dr.

Rudolf Meier (MfN), Dr. Paula Riccardi (MfN) and Vivian Yang (MfN) for helpful input and commentary on this program while it was being coded.

## Appendices

### Appendix 1. List of Sclerites Used

entire_forewing, pterostigma, cell-csc, cell-Rs, cell-cu1, cell-cu2, cell-m1, cell-m2, cell-r2, cell-a

### Appendix 2. Example taxonomic species description snippet output from *Descriptron*

Species Description for *nana*:

### Material Examined

- **Locality**: Kapama Game Reserve; **Scientificname**: Sclerocarya birrea caffra;

**Family**: Anacardiaceae; **Country**: South Africa; **Stateprovince**: Limpopo;

**County**: Mopani District Municipality; **Locality**: Kapama Game Reserve;

**Decimallatitude**: -24.403; **Decimallongitude**: 30.994;

**Coordinateuncertaintyinmeters**: None; **Elevation**: 550; **Elevationaccuracy**: None;

**Eventdate**: 2021-09-08; **Recordedby**: D. Burck

### Description

The forewing of *nana* is broad with a short width-to-length ratio and an oblong oval shape. It features pale brown veins and a completely hyaline membrane, moderately covered with spinules forming a pattern of two hexagonal rows. The veins are pale yellow, with no color pigmentation present along the apices of the pterostigma and veins Rs, M1+2, M3+4, Cu1a, and Cu1b.

Spinule-free strips are narrow and present along all veins. The apical two-thirds of vein C+Sc is nearly straight, and there is no costal break between vein C+Sc and the pterostigma. The pterostigma cell is absent. Vein Rs is relatively straight and slightly turns up to the wing margin apically. Cell M1 is broad and short, while cell Cu1 is broad and long. Vein Cu1a is strongly convex along its entire length, with its apex moderately arced towards the posterior margin. The forewing is widest in the middle, with a broadly and evenly rounded apical margin, and is without setae.

### Morphological Measurements

- **Entire Forewing**: Length 1.2295 mm, Height 0.4840 mm, Area2 0.4466 mm, Perimeter 3.0114 mm, Equivalent diameter 0.7541 mm
- **Cell-m1**: Length 0.3713 mm, Height 0.1719 mm, Area2 0.0406 mm, Perimeter 0.9839 mm, Equivalent diameter 0.2274 mm
- **Cell-m2**: Length 0.5857 mm, Height 0.1258 mm, Area2 0.0592 mm, Perimeter 1.4086 mm, Equivalent diameter 0.2745 mm
- **Cell-r2**: Length 0.9319 mm, Height 0.1428 mm, Area2 0.0864 mm, Perimeter 2.1418 mm, Equivalent diameter 0.3317 mm
- **Cell-cu2**: Length 0.6019 mm, Height 0.1875 mm, Area2 0.0902 mm, Perimeter 1.4719 mm, Equivalent diameter 0.3390 mm
- **Cell-cu1**: Length 0.3538 mm, Height 0.1225 mm, Area2 0.0341 mm, Perimeter 0.8669 mm, Equivalent diameter 0.2084 mm
- **Cell-Rs**: Length 0.6584 mm, Height 0.1073 mm, Area2 0.0600 mm, Perimeter 1.4675 mm, Equivalent diameter 0.2763 mm
- **Pterostigma**: Length 0.4565 mm, Height 0.0381 mm, Area2 0.0109 mm, Perimeter 0.9698 mm, Equivalent diameter 0.1176 mm

### Diagnosis

*nana* is distinguished by its broad forewing with a short width-to-length ratio, pale brown veins, and a hyaline membrane. The absence of a pterostigma cell and the presence of a moderately arced apex of vein Cu1a are key features that differentiate it from other species.

……………………….. Dichotomous Key:

1a. Forewing pale yellow go to 2

1b. Forewing not pale yellow go to 8

2a. Forewing with transverse row spinule pattern *fabianae*

2b. Forewing with different spinule pattern go to 3

3a. Forewing with randomly distributed spinules *similis*

3b. Forewing with different spinule arrangement go to 4

4a. Forewing with costal break *chorizanthis*

4b. Forewing without costal break go to 5

5a. Forewing with narrow spinule-free strips *bicolorata*

5b. Forewing with broad spinule-free strips go to 6

6a. Forewing with dark brown pigmentation along pterostigma *mendozae*

6b. Forewing without dark brown pigmentation go to 7

7a. Forewing with distinct brown patches *longicauda*

7b. Forewing without distinct brown patches *similis*

8a. Forewing pale brown go to 9

8b. Forewing not pale brown go to 15

9a. Forewing with pale brown veins go to 10

9b. Forewing with different vein coloration go to 12

10a. Forewing with hyaline membrane *nana*

10b. Forewing without hyaline membrane go to 11

11a. Forewing with transverse rows of spinules *adela*

11b. Forewing with different spinule arrangement *punctulata*

12a. Forewing with two hexagonal rows of spinules go to 13

12b. Forewing with different spinule arrangement *theresae*

13a. Forewing with short, densely distributed setae *nolanae_female*

13b. Forewing without setae *brevigenis*

14a. Forewing with broad spinule-free strips *dimorpha_female*

14b. Forewing with narrow spinule-free strips *lycii_female*

15a. Forewing oblong oval go to 16

15b. Forewing not oblong oval go to 18

16a. Forewing with pale brown color *viscosae*

16b. Forewing with different color pattern go to 17

17a. Forewing with dark brown veins *magellanica*

17b. Forewing with pale yellowish-brown color *longirostro*

18a. Forewing narrow and long *sebastiani*

18b. Forewing broad and short *lycii_male*

19a. Forewing with distinct brown veins and patches *adunca*

19b. Forewing with different vein and patch pattern *theresae*

---

**SUPPLEMENTAL FIGURE S1.**
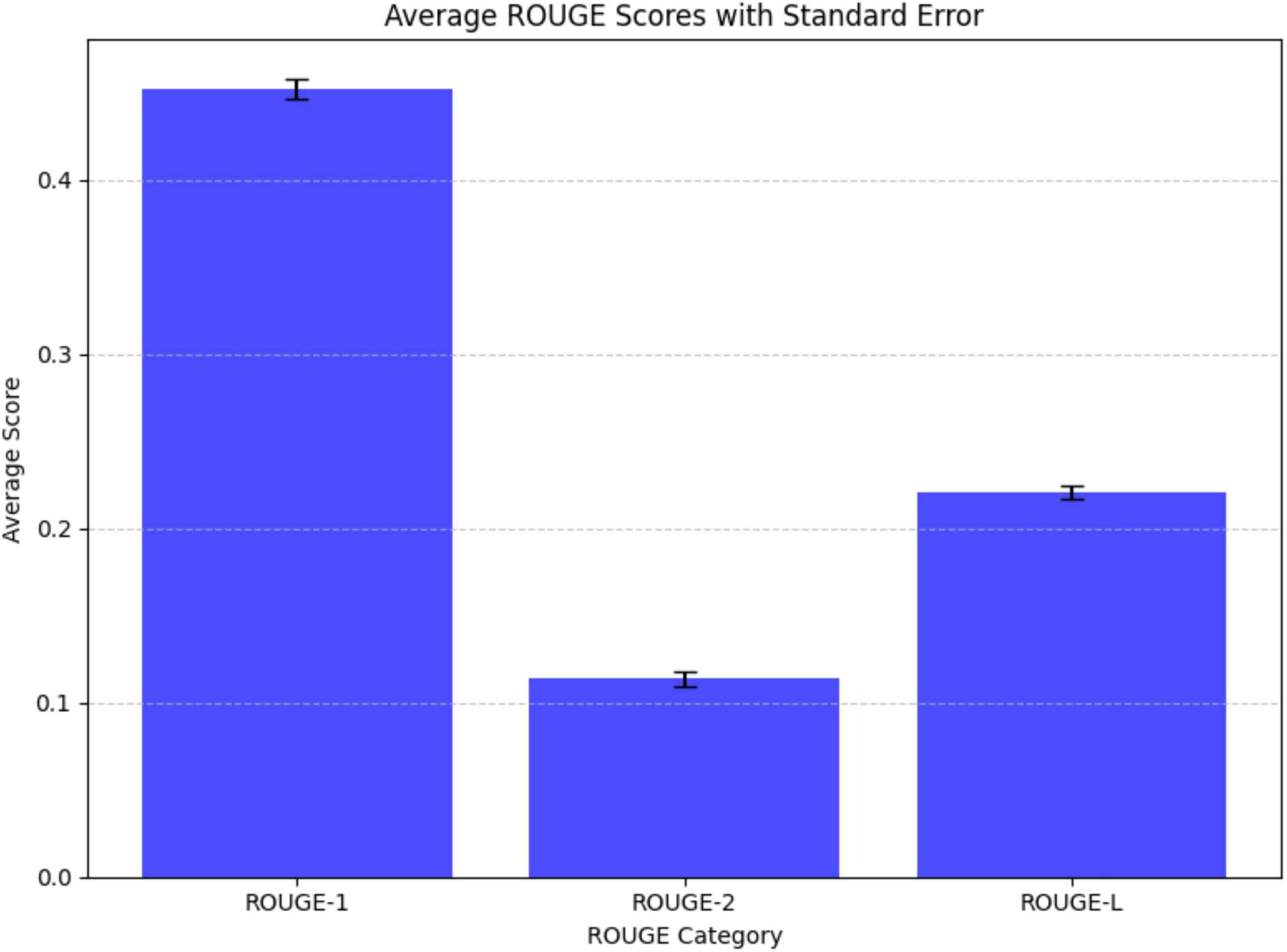
Average ROUGE scores with standard error bars comparing taxonomist and GPT-4o model output from 43 *Russelliana* specimens examined.

**SUPPLEMENTAL FIGURE S2.**
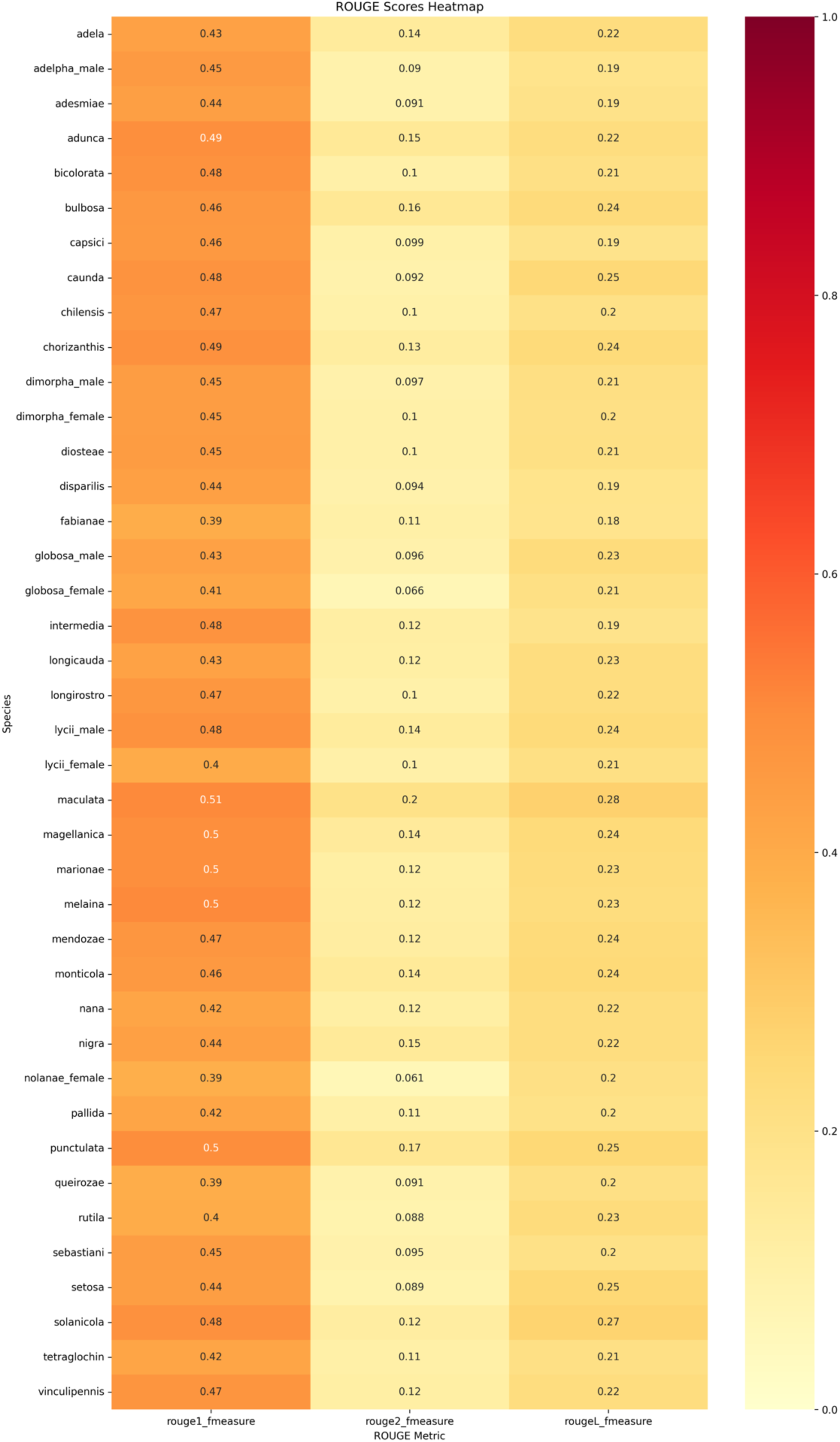
Rouge scores from 43 *Russelliana* samples comparing taxonomist and GPT-4o species forewing descriptions.

**SUPPLEMENTAL FIGURE S3.**
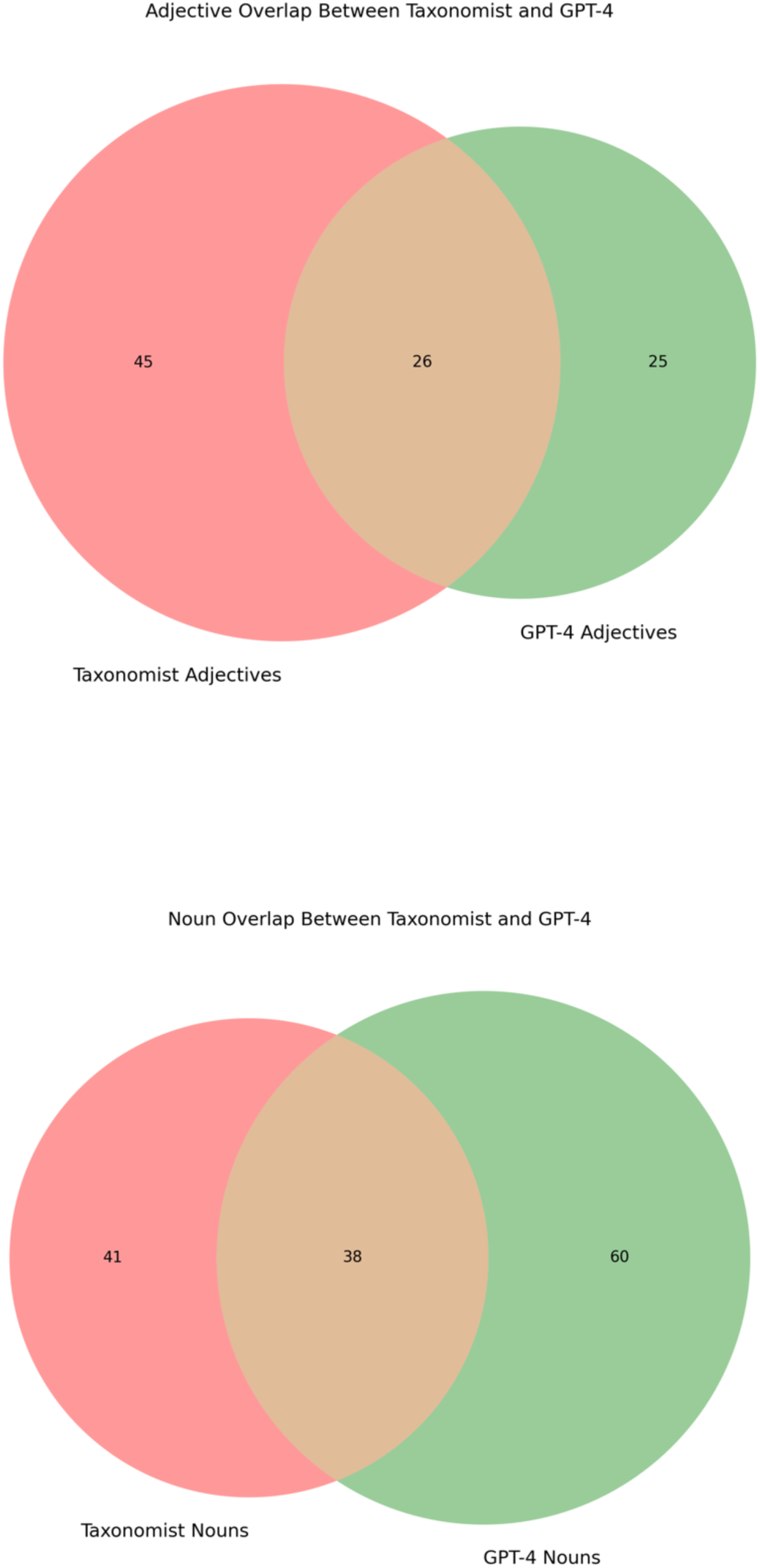
Venn diagrams comparing Adjective and Noun overlap between Taxonomist and GPT-4o forewing species descriptions from 43 *Russelliana* specimens.

**SUPPLEMENTAL FIGURE S4.**
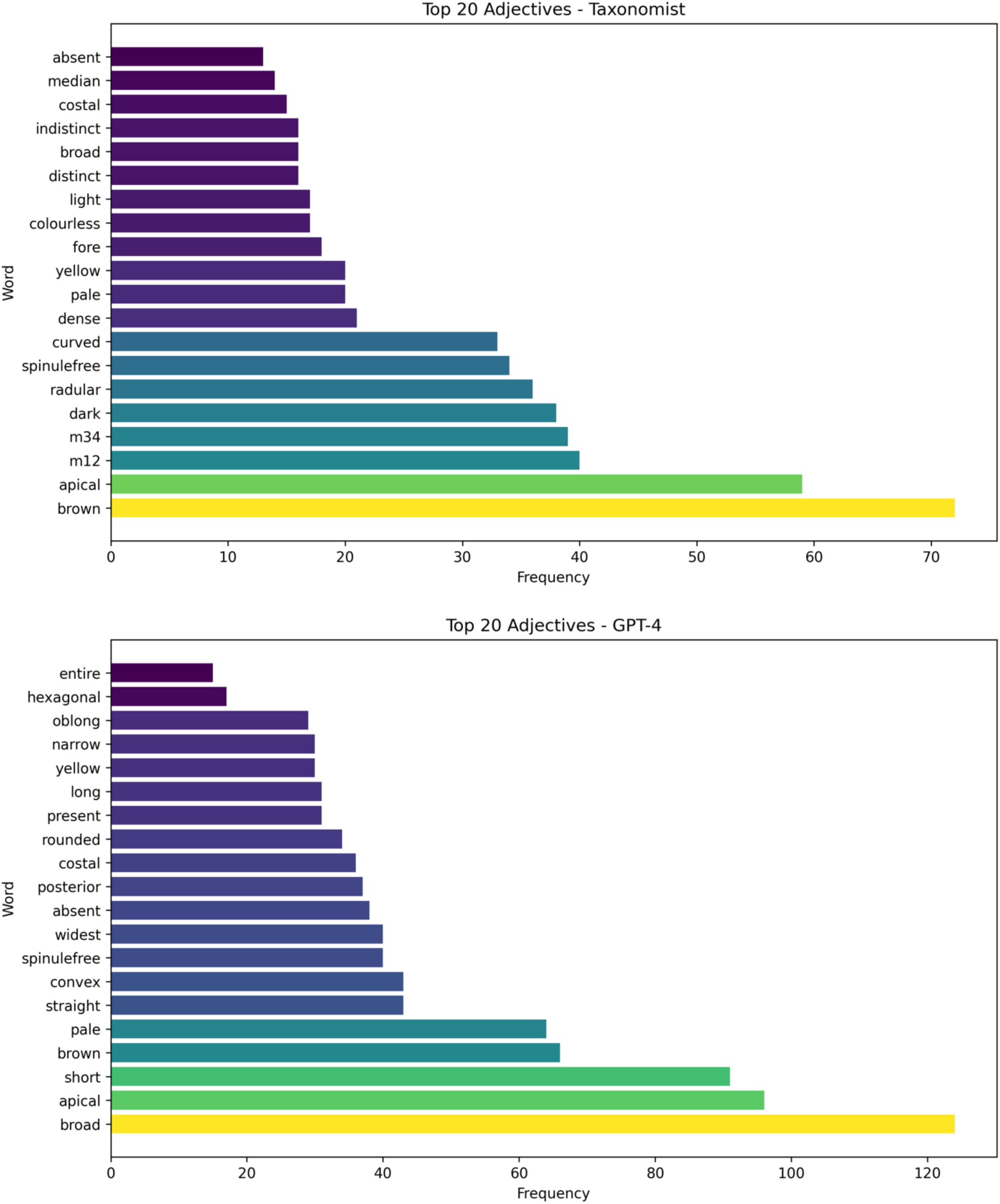
Taxonomist and GPT-4o top 20 most common Adjectives used in forewing descriptions of 43 *Russelliana* specimens.

**SUPPLEMENTAL FIGURE S5.**
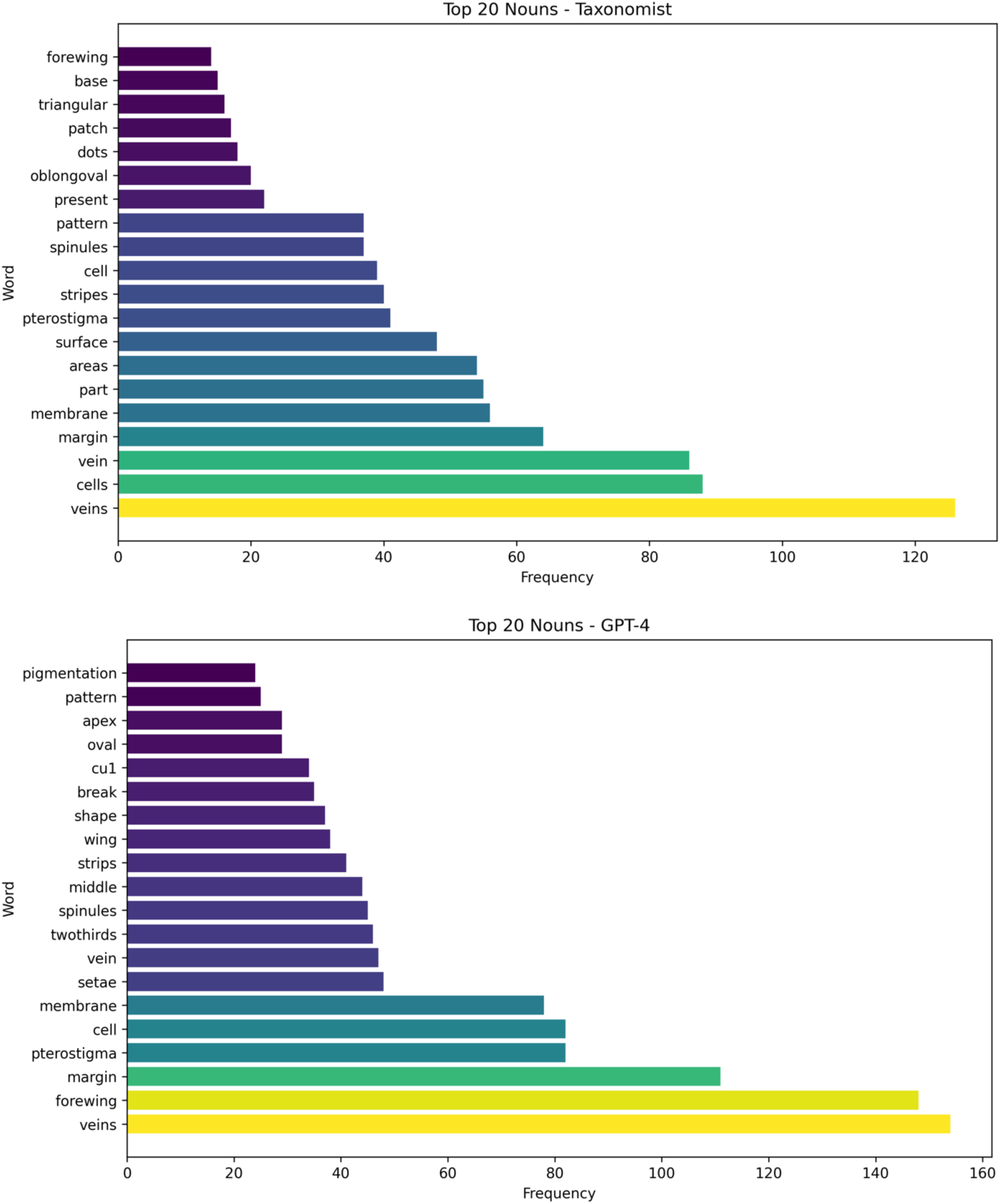
Taxonomist and GPT-4o top 20 most common Nouns used in forewing descriptions of 43 *Russelliana* specimens.

**SUPPLEMENTAL FIGURE S6.**
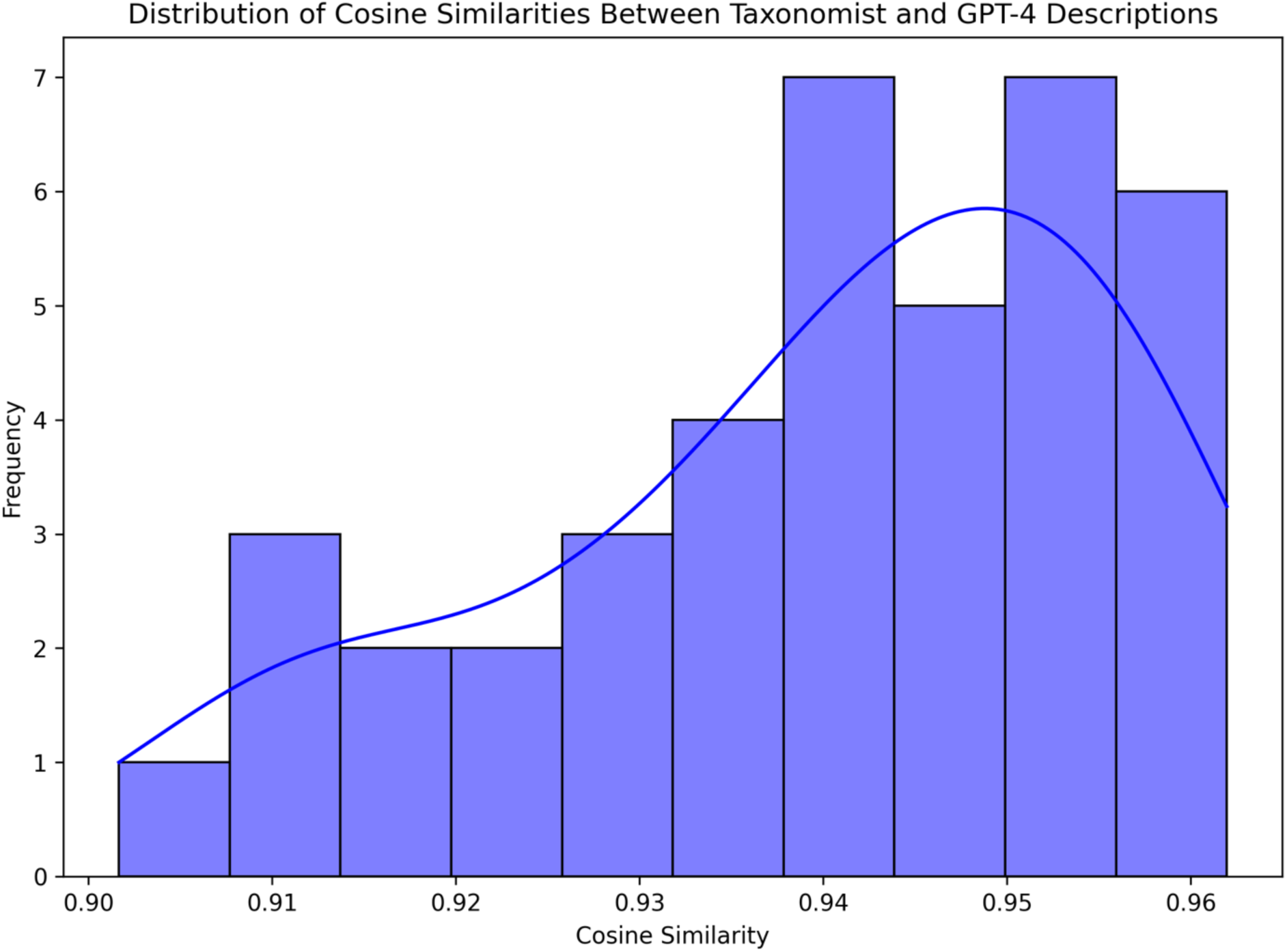
Cosine similarity scores comparing forewing descriptions between 43 *Russelliana* specimens generated by taxonomists and GPT-4o.

